# Multi-modal Ensemble Approach for Decoding Player Intentions in Table Tennis

**DOI:** 10.64898/2026.05.04.721564

**Authors:** Trung Quang Pham, Shotaro Shiba Funai, Ryota Kanai, Junichi Chikazoe

## Abstract

This study aims to predict human intentions during intense sports activities, specifically in table tennis. Using a publicly available Real World Table Tennis dataset containing simultaneous EEG and video recordings, we developed a series of subject-specific classifiers for nine players (7 males and 2 females; age range 18–30), based on pose features and EEG signals. The pose-based classifier used a stochastic gradient descent model with logistic loss, whereas the EEG-based classifier employed a modified convolutional neural network architecture (EEGNet). Both classifiers successfully predicted left-right attack intentions from the time windows preceding racket-ball impact, with optimal decoding occurring at −100 ms for pose features and −500 ms for EEG signals. Although EEG-based decoding achieved comparable performance to pose-based decoding, a multi-modal ensemble further improved prediction significantly, reaching a mean macro F1 score of 0.563 (bootstrapped 95% CI: 0.523–0.603), corresponding to gains of +0.03 over pose-only and +0.02 over EEG-only classifiers. Because each classifier is trained independently, the ensemble can be feasibly extended to incorporate additional modalities in the future. These results suggest potential relevance of this framework as neurofeedback tools for sports training and neural prosthetic systems in future work.

## Introduction

Intention decoding, particularly through noninvasive methods, has been drawing attention in the brain-machine interface (BMI) field in recent years. BMI fills the gap between brain activity and physical movements. Therefore, an effortless and accurate intention decoding is crucial for the development of a system such as a neural prosthetic^1,2^ or brain-to-text handwriting^3^.

Various techniques have been developed to decode the intention of movement using electroencephalography (EEG)^4–8^. Early approaches typically relied on hand-crafted feature extraction methods, such as common spatial patterns, followed by conventional machine learning classifiers^9–12^, whereas more recent work has adopted deep learning models that operate directly on raw EEG signals^11,13–20^. While these methods have shown competitive performance under controlled conditions, EEG’s susceptibility to motion artifacts has led most studies to focus on the imagination of movement, which is slow and may not accurately reflect the actual action planning, particularly in sports. In intensely competitive sports such as table tennis, movement could occur within a few hundred milliseconds and often relies on the subconscious mind. Recent advances in EEG technology have led to the development of portable headsets, enabling brain recording during movement-intensive activities such as walking, cycling^21–23^, and even playing table tennis^24^. These hardware achievements have opened up a more innovative and dynamic approach to intention decoding.

Table tennis is a globally popular sport and has been part of the Summer Olympic Games since 1988. Due to its demand for full-body coordination, precise timing, and rapid-decision making, table tennis has been widely adopted in both behavioral studies^25,26^ and neurophysiological studies^27,28^. This makes table tennis a good candidate for investigating intention decoding in dynamic, real-world scenarios.

Traditionally, detecting the intention that underlies an attack has relied on analyzing the trajectory of the table tennis balls (TBBs)^29–33^. However, TBB trajectories are influenced by numerous physical factors, making accurate detection both complex and time-consuming. A more direct and reliable indicator would be the player’s pose^34,35^ in response to an incoming ball. Nevertheless, in professional matches, players often employ deceptive movements as part of their strategy, making pose-based intention decoding challenging.

With regard to intentions, brain activity may serve as an ideal indicator^36^. In highly trained players, neural responses occur more instinctively and are therefore less consciously alterable. A study involving professional, young elite, and amateur players found a relationship between skill level and motor cortex excitability during reaction, movement planning, and execution under high attention demands^37^. Additional research has shown strong associations between the table tennis professionalism and enhanced neural efficiency in visual processing areas, as well as faster visuomotor reaction times^38,39^. Furthermore, magnetic resonance imaging (MRI) studies have revealed stronger activation in experts—compared to novices—in the semantic and sensorimotor areas, suggesting that long-term table tennis training may improve dynamic functional connectivity in large-scale brain regions^27,40^.

In the domain of brain modeling, there is no universally dominant model. Instead, a recent theory of intelligence—known as “The Thousand Brain Models”^41^—proposes that the brain prefers to learn multiple models for a concept rather than relying on a single one. Despite learning from different inputs, the models eventually vote to reach a consensus of what they are recognizing. This theory is analogous to the concept of “multi-modal ensemble learning” in machine learning. This approach has been shown to improve performance across a wide range of tasks, including zero-shot image recognition^42^, emotion recognition^43^, language translation^44^, action recognition^45^, as well as complex medical tasks^46–48^. In many domains, “modalities” often refer to different types of physiological data (e.g., functional MRI, EEG, EMG) which primarily capture the internal states. However, pose information, which reflects external physical states through joint positions, joint angles, can also be considered as a video-based kinematic modality. Therefore, the combination of multiple modalities, such as neural modality (EEG) and kinematic modality (pose), has strong potential to enhance both the accuracy and robustness of intention decoding systems.

However, most existing intention decoding studies rely on a single modality and are evaluated under motor imagery or highly controlled experimental paradigms. Decoding intentions during fast, continuous, and physically demanding real-world tasks, such as competitive sports, therefore remains underexplored. As a result, it remains unclear whether or how internal neural signals and external kinematic cues can be combined in a principled manner to improve the robustness of intention decoding in realistic settings.

To bridge this gap, we investigate the efficacy of integrating kinematic pose data with electroencephalography (EEG) signals to decode player intentions during dynamic, real-world table tennis matches. Focusing on foundational left-versus-right attack intentions, we propose a modular multimodal ensemble framework that combines independently optimized, subject-specific EEG and pose classifiers. Using synchronous brain activity and video footage from a publicly available dataset^24^, our results demonstrate that integrating neural and kinematic modalities significantly enhances intention-prediction accuracy in competitive sporting environments.

The main contributions of this study are as follows:

- We demonstrate that motor intentions can be decoded 100 to 500 ms prior to racket–ball impact without requiring explicit motor imagery or specialized pre-training. Given that table tennis execution phases typically span less than 100 ms^49^, predictions within this window reflect early neural and kinematic signatures of motor planning and preparation^8,50–52^.
- We introduce novel kinematic features for pose-based classifiers, and a transfer-learning approach for EEG-based classifiers. Both modifications substantially improve the performance of their respective models.
- We propose a modular multi-modal ensemble framework that integrates independently trained neural (EEG) and kinematic (pose) classifiers, allowing complementary modality-specific strengths to be leveraged while maintaining scalability and extensibility for broader neurotechnology applications and sports analytics.

The rest of this paper is organized as follows. The “Literature review” section provides a comprehensive background of related studies. The “Methods” section describes our proposed framework, the dataset used, the participant selection process, data labeling, implementation details, and analysis procedures. The “Results” section provides the development of the pose-based and EEG-based classifiers, the proposed ensemble approach, and the corresponding evaluation results. The “Discussion” section contextualizes our findings and addresses study limitations. Finally, the “Conclusion” section summarizes the results and highlights the novelty of this current work.

## Literature review

### Intention inference from kinematics and pose in sports contexts

Intention decoding using either pose information or EEG signals has attracted increasing attention in recent years. Cavallo et al.^53^ demonstrated that humans can infer intention-related information from movement kinematics. Although their study focused on the grasping movements, this finding has motivated broader investigations into the relationship between kinematic features and human intention^54,55^. Extending to full-body movement based prediction, Gesnouin et al.^56^ showed that pedestrian intention can be decoded from 2D pose sequences using deep neural networks. However, intention decoding based on movement kinematics remains largely unexplored in sports settings, particularly table tennis. Related foundational work includes Wu et al.^34^, who implemented a real-time pose estimation for Table Tennis using the OpenPose model^57^, and Wang et al.^35^ who provided a comprehensive tennis player action dataset that may support future developments in intention prediction.

### EEG-based intention decoding beyond motor imagery

For intention decoding from neural activity, most studies have focused on “motor imagery” tasks, in which participants are instructed to imagine their movements. This is partly due to the availability of large, well-established public datasets, such as the EEGMMIDB dataset^58^, BCI Competition III^59^, BCI Competition IV^60^. Traditional approaches rely heavily on hand-crafted feature extraction methods, most notably the Common Spatial Pattern (CSP) method and its variants, followed by simple classifiers, including linear discriminant analysis (LDA)^9^, random forest^10^, support vector machine (SVM)^11^, and extreme learning machine (ELM)^12^. Previous studies have also examined the influence of classifier choice and feature representation, showing that performance varies depending on kernel selection, feature dimensionality, and subject variability, particularly for SVM- and ELM-based approaches^61^. This leads to substantial differences across datasets and experimental settings.

Beyond EEG, neural intention decoding has also been examined using modalities such as fMRI. Gallivan et al.^62^ successfully decoded the reach-and-grasp intentions from fMRI signals in the parieto-frontal networks, whereas Ruiz et al.^51^ extended this to motor imagery and highlighted the involvement of additional areas, such as the premotor cortex. These insights would be useful for selectively reducing the redundant electrodes in EEG recordings, enabling more comfortable and practical recording sessions.

To address electrode redundancy, Li et al.^63^ introduced the Gradient-Weighted Class Activation Mapping (Grad-CAM) visualization technology alongside a recurrent convolution neural network model. Their method reduces the number of electrodes by half while maintaining relatively high accuracy on the EEGMMIDB dataset^58^. However, since reduced-channel performance remains noticeably lower than that of the full-channel model, the trade-off between model performance and channel number must be considered. For complex tasks (such as in this study) where performance is critical, retaining all channels may be necessary.

Compared to pre-trained motor imagery tasks, real-world motor tasks are inherently more complex and noisy^7^. To address the limitations of traditional motor-imagery-based approaches, Ganesh et al.^8^ introduced novel sensory prediction errors which allow for predicting whether wheelchair users thought of turning left or right. Using galvanic vestibular stimulation (GVS) as feedback, they achieved 87.2% accuracy without any user training. Although GVS has been investigated for sports applications, its post-stimulation effects may be unsuitable for high-intensity sports like table tennis.

A notable study by Salvaris et al.^5^ further challenged the assumption that repetitive motor imagery is mandatory. They used a combination of sensorimotor rhythms and motor imagery training for successfully decoding left- and right-hand movement intentions at sub-second timescales. Although free-choice actions were decoded for a subset of high-performing participants, the requirement for motor imagery training and the limited generalization present challenges for applying similar approaches to real-world motor tasks, particularly in sports.

### Lightweight EEG deep models and transfer learning

The inherent limitations of the traditional approaches have motivated recent efforts toward deep learning approaches that aim to learn task-relevant representations directly from raw or minimally processed signals, reducing dependence on manual feature engineering. Lawhern et al.^13^ introduced the EEGNet architecture, which achieved state-of-the-art performance across multiple paradigms, including movement-related cortical potentials and sensorimotor rhythms. Following this, various deep neural networks have been applied for motor imagery tasks, such as convolutional neural networks (CNNs)^14,15^, recurrent neural networks (RNNs)^16^, long-short-term memory (LSTM)^17^, Transformers^18^, and hybrid convolutional Transformer networks^19,20^. Despite these developments, EEGNet remains the preferred baseline due to its lightweight structure, ease of modification, and suitability for real-time application^64^. Notably, Wang et al.^15^ demonstrated that the EEGNet can even be deployed on low-power microcontroller units with only a minimal accuracy reduction of 0.32%.

### Multi-modal and ensemble approaches for intention and behavioral decoding

Recent advances in decoding complex human behavior have leveraged multi-modal and ensemble learning frameworks to improve prediction performance. For example, hybrid ensemble models have been successfully applied to human activity recognition tasks^45^, demonstrating the potential of combining diverse feature representations. In the specific context of motor expertise and action representation, studies have shown that expertise shapes cross-modal and modality-specific representations in table tennis players, highlighting the importance of integrating kinematic and neural information for understanding intention and action^65^. These developments suggest that multi-modal ensemble approaches which leverage complementary strengths of neural and kinematic modalities have strong potential for improved intention decoding in real-world tasks.

This literature review indicates that most existing studies on intention decoding rely on a single modality, such as EEG or kinematic information (Table 1). Earlier approaches primarily depended on handcrafted feature extraction, whereas recent work has increasingly adopted deep learning–based models. Although these methods often achieve high classification accuracy, many are evaluated under motor imagery or cue-based experimental paradigms, which differ substantially from continuous, naturalistic tasks. In contrast, our study focuses on decoding player intentions from EEG and pose signals recorded during a continuous, real-world table tennis task, using offline analysis of segmented data and without requiring explicit motor imagery. By leveraging a modular decision-level ensemble, our approach exploits complementary neural and kinematic information while maintaining robustness and interpretability in movement-intensive conditions. This distinguishes our work from prior single-modality or imagery-based intention decoding studies.

**Table 1.** Qualitative comparison of representative studies on intention decoding and related tasks, highlighting differences in modality, task focus, experimental setting, prediction timing, and fusion strategy. N/A: Not applicable.

| Study | Modality | Task focus | Real-world settings | Prediction timing | Multi-modal fusion | Year |
| --- | --- | --- | --- | --- | --- | --- |
| Gesnouin et al.<sup>56</sup> | Pose | Intention decoding | Yes | Pre-action | No | 2020 |
| Wu et al.<sup>34</sup> | Pose | Pose estimation | Yes | N/A | No | 2023 |
| Wang et al.<sup>35</sup> | Pose | Pose estimation | Yes | N/A | No | 2024 |
| Gaur et al.<sup>9</sup> | EEG | Motor imagery | No | Cue-based | No | 2021 |
| Sharma et al.<sup>18</sup> | EEG | Motor imagery | No | Cue-based | No | 2023 |
| Song et al.<sup>19</sup> | EEG | Motor imagery | No | Cue-based | No | 2023 |
| Zhao et al.<sup>20</sup> | EEG | Motor imagery | No | Cue-based | No | 2024 |
| Salvaris et al.<sup>5</sup> | EEG | Intention decoding | Yes | Pre-action | No | 2014 |
| Ganesh et al.<sup>8</sup> | EEG | Intention decoding | Yes | Pre-action | No | 2018 |
| Li et al.<sup>63</sup> | EEG | Intention decoding | No | Pre-action | No | 2020 |
| Gallivan et al.<sup>62</sup> | fMRI | Intention decoding | No | Pre-action | No | 2011 |
| Ruiz et al.<sup>51</sup> | fMRI | Intention decoding | No | Pre-action | No | 2024 |
| Topuz et al.<sup>45</sup> | Biometrics | Activity recognition | Yes | Post-action | Yes | 2025 |
| **Ours** | **EEG + Pose** | **Intention decoding** | **Yes** | **Pre-impact** | **Yes (ensemble)** |  |

## Methods

### Dataset

We used the Real World Table Tennis dataset^24^, which was collected and published by a separate research group. The dataset includes synchronized EEG recordings and full-body video data acquired during gameplay, along with survey responses. All participants are right-hand dominant with normal or corrected-to-normal vision and no musculoskeletal or neurobiological injuries. Most participants casually play with family and friends (ranging from 1 to 500 games).

Among 25 subjects in the dataset, 9 subjects (7 males and 2 females; age range 18–30; Supplementary Table S1) were selected for this study. The selection criterion was primarily based on their responses related to playing frequency (Q1), tournament history (Q2, Q6), self-assessment (Q3, Q7). As Q1, Q2, and Q3 were directly related to table tennis, while Q6 and Q7 were related to other racquet sports, responses to Q1–Q3 and Q9 were prioritized. Based on this criterion, 12 subjects with low scores in both Q1 and Q2 were excluded, specifically those who reported playing table tennis less than once per month and having never attended an organized event. Additionally, four subjects were excluded due to data quality issues. Two subjects (sub-01, sub-18) were excluded due to the lack of corresponding video recordings. Two subjects (sub-10, sub-25) were excluded because the total number of left and right attacks was below 400, which is insufficient for robust training and evaluation. Participant’s survey responses, data availability, and our inclusion decision are shown in Supplementary Table S2.

The rasket-ball impact frames were extracted and manually labeled using the markers in the corresponding Adobe Premiere Pro files. Three labels were used: left, right, and in-between, encoded as 0, 1, and 2 respectively. To ensure the analyzed data reflected deliberate intent rather than random or purely reflexive actions, only trial sequences in which subjects exhibited a clear, stable, and well-defined movement trajectory (judged by an external observer) were retained for subsequent analysis.

EEG signals were recorded using a multi-channel EEG system, while body pose information was obtained using a markerless motion capture approach. The experimental protocol, sensor specifications, acquisition parameters, and ethical approvals are described in detail in the original publication^24^. In the present study, no new data collection was performed; all analyses were conducted on the preprocessed EEG signals released with the dataset.

### Multi-modal Ensemble Method

Figure 1 illustrates our proposed multi-modal framework. EEG signals and kinematic (pose) information are processed through two parallel, modality-specific pipelines, each consisting of a dedicated preprocessing or feature extraction stage followed by an independent classifier. The outputs of these classifiers are then combined using a decision-level ensemble model to produce the final prediction.

**Figure 1.**
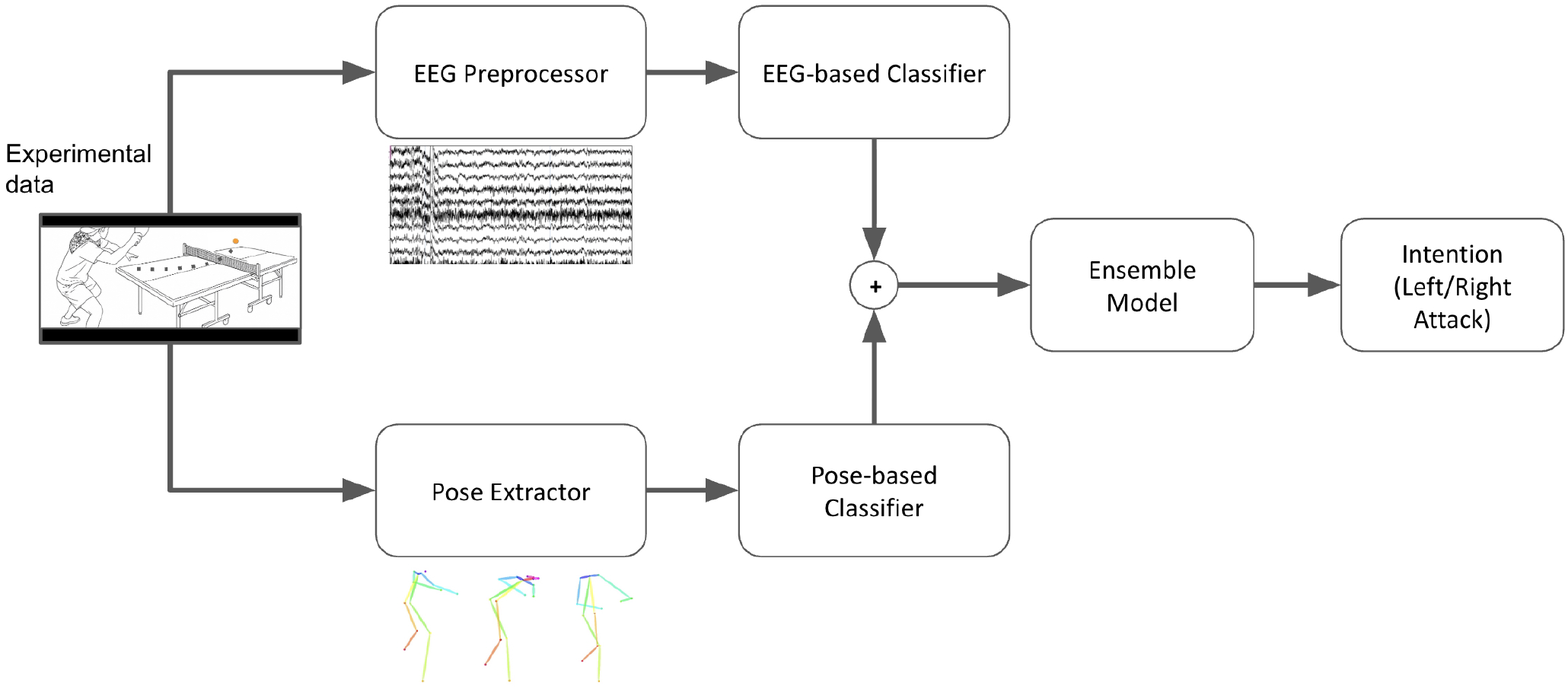
Schematic overview of the proposed multi-modal ensemble framework. EEG and pose features are processed in parallel and combined via an ensemble model to predict left–right attack intentions.

This modular architecture allows each modality to be modeled and optimized independently, which is particularly advantageous in movement-intensive scenarios where signal quality and noise characteristics differ substantially between EEG and pose data. While more tightly coupled feature-level fusion strategies have been explored in other domains^66,67^, we intentionally prioritize decision-level integration to preserve modality-specific representations. Moreover, the decision-level fusion strategy enables flexible scaling of classifiers within each modality while maintaining the robustness and interpretability of the overall system.

### Preprocessing of EEG data

According to the original dataset documentation, the data were processed using a 1 Hz high-pass filter to remove slow drifts, 60 Hz line-noise removal, low-pass filtering at 125 Hz, and downsampling to 250 Hz, together with motion- and muscle-artifact removal. Bad channels (defined as those exceeding three standard deviations from the median voltage across channels) were rejected and subsequently interpolated using spherical interpolation. To ensure full-rank data, principal component analysis (PCA) was applied by the dataset authors, followed by adaptive mixture independent component analysis (AMICA ^68^). In this study, we used these preprocessed EEG signals directly, and did not apply additional preprocessing steps beyond excluding non-EEG channels.

### Temporal Segmentation and Modality Synchronization

Session boundaries were defined based on the associated event files. In total, 16 sessions were analyzed, each corresponding to a specific video recording. Start and end times for each session were determined from the recorded trial types and the first subject hit, as detected by the paddle-mounted inertial measurement unit (IMU) accelerometer. In the video data, all hitting markers were time-stamped relative to this first hit event. Accordingly, we used the first hit of each session to temporally synchronize the video and EEG data. The number of subject-hit events was cross-checked against the number of hitting markers in the video recordings. Additionally, we computed the intervals between consecutive hits in both the EEG and video data to verify precise temporal alignment.

### ERS/ERD Analysis

Event-related synchronization/de-synchronization (ERS/ERD) analysis was conducted as an auxiliary visualization to examine whether canonical motor-related spectral patterns were observable in the pre-impact interval (Supplementary Fig. S1). ERS/ERD was computed as:

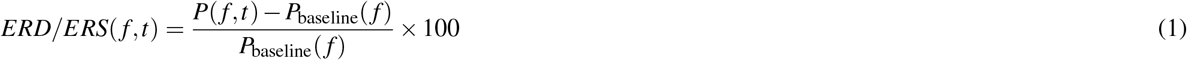

where *P*(*f* , *t*) is the time–frequency power at frequency *f* and time *t*, and *P*_baseline_ is the mean power during the baseline interval (− 2.0 to −1.5 s relative to impact). Negative values indicate ERD, while positive values indicate ERS.

ERD/ERS time courses were first computed at the subject level (using MNE-Python library v1.6.0^69^), and interpolated to a common time grid. Group-level averages and 95% confidence intervals (CIs) were then calculated across subjects for the *µ* (8–13 Hz) and *β* (13–30 Hz) frequency bands at electrodes C3 (left) and C4 (right). This analysis was not used for classification or model optimization.

### Pose extraction

To extract the pose data from video clips, we used the OpenPose (v1.0.0) model^57^. The pose information at −100 ms before impact was extracted for each hit. This timing was selected based on a preliminary model evaluation at multiple time points (0 s, − 100 ms, − 200 ms, −500 ms, −1 s, and −2 s before impact).

The original pose output includes keypoints for the arms, elbows, body, neck, head, and legs. Head-related keypoints were excluded due to the censorship in the video clips. From the remaining keypoints, we derived pose features by computing the coordinates of connected links based on paired points. In addition, we calculated the orientation (*θ*) and length (L) for each link as follows,

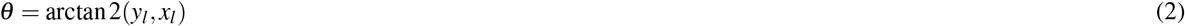

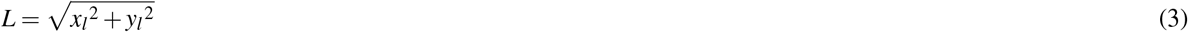

where *x*_*l*_, *y*_*l*_ are the horizontal and vertical components of the *l*^*th*^ link, respectively.

### Classifier preparation

Several classifiers were built to predict the player intention. To keep the analysis straightforward, we focus on the left-right attack intentions, thus framing the problem as a typical binary (two-classes) classification task. The class-wise sample counts after data selection are provided in Supplementary Table S1.

A pose-based classifier was implemented using a stochastic gradient descent (SGD) classifier with logistic loss and balanced class weights. We applied ElasticNet regularization, where the regularization strength (*α*) was selected through cross-validation. The objective function is given as follows:

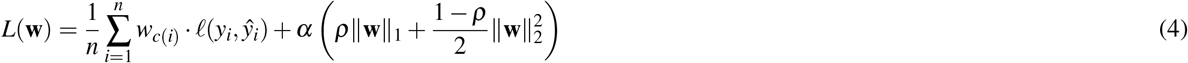

where *n* is the number of samples, *w*_*c*(*i*)_ is the class weight for the *i*^*th*^sample, *ℓ*(*y*_*i*_, *ŷ*_*i*_) is the logistic loss, **w** is the feature weight vector, *α* is a multiplies constant for the regularization term, *ρ* is the norm *L*_1_ ratio, a mixed regularization value for the norms *L*_1_ and *L*_2_. The *α* parameter was tested in logarithmic steps from 10^−10^ to 10^4^. We used the SGD classifier implementation in scikit-learn library v1.0.2^70^.

The EEG-based classifiers were built on the EEGNet architecture^13^. Although many newer CNN- and Transformer-based models have been proposed^16–20^, EEGNet remains a standard baseline in EEG-based studies^64^, demonstrating a robust and stable performance across various tasks, including motor imagery classifications and stroke states classification, even when trained on relatively small datasets. Furthermore, EEGNet is sufficiently lightweight to support real-time application, making it well-suited for future online implementations^15,64^. Given the size of the dataset used in this study, we selected EEGNet for its compact yet efficient architecture, based entirely on convolutional layers, which facilitates straightforward modification and adaptation.

For architectural validation, we additionally implemented several baseline EEG classifiers, including an SGD model with ElasticNet regularization (*α* ranging from 10^−10^ to 10^4^ in logarithmic steps) and two Transformer-based architectures (EEGConformer^19^ and CTNet^20^). These models were trained and evaluated using the same subject-specific cross-validation protocol as EEGNet. For the Transformer-based models, we adopted the data augmentation strategy proposed by Song et al.^19^ to mitigate the impact of limited sample sizes.

We modified the input layer to accommodate all 120 EEG channels in our dataset and adjusted the output layer to perform binary classification of attack direction (0 for left, 1 for right). The model parameters are shown in Supplementary Fig. S2. The dropout probability was set at 0.25. EEG signals were used without any additional feature extraction.

The modified EEGNet was trained using two distinct approaches:

1. **Direct Training Approach**
2. **Transfer Learning Approach**

In the first approach, we trained the EEGNet directly on each subject’s data for two distinct binary classification tasks (Left vs. Others and Right vs. Others). The intermediate (in-between) trials were assigned to the ‘Others’ class during training; this design choice served as a regularizer, forcing the classifier to learn sharper decision boundaries for the primary target classes. The input was randomly divided into a training set and a validation set, and early stopping was applied if the validation loss failed to improve after 50 epochs. We used the Adam optimizer^71^ with hyperparameters such as learning rate, batch size, and weight decay tuned via 8-fold cross-validation. Due to class imbalance (a predominance of left attacks), we calculated class weights based on the training data and applied random oversampling using the imbalanced-learn Python package^72^.

The loss function used was weighted cross-entropy, defined as follows:

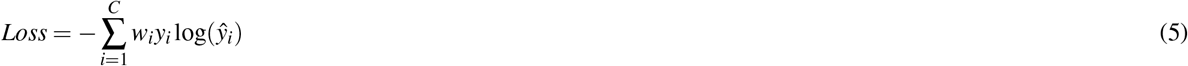

where *w*_*i*_ is the pre-computed weight assigned to class *i, y*_*i*_ is the ground-truth binary indicator (0 or 1) for class *i*, and ŷ_*i*_ is the predicted probability (softmax output) for class *i*, and C is the number of classes. It should be noted that while binary classification is commonly implemented using a single output neuron with a sigmoid activation, our implementation of two output neurons with a softmax activation yields identical decision boundaries under cross-entropy loss. A proof of equivalence is provided in the Supplementary Information.

In the second approach, we leveraged a transfer learning technique to optimize the initialization of the shallow layers. This approach is motivated by the limited size of each individual dataset. A generalized group model was pre-trained using a leave-one-subject-out (LOSO) strategy (training on all subjects except the target subject). Pre-training on group data is expected to improve the initialization of the shallow layers, thereby enhancing generalization when fine-tuning on subject-specific data. Similar methodologies have been proposed by Pham et al.^73^ and Hasson et al.^6^. Adam optimizer was used with a learning rate of 0.0001 and a weight decay of 0.00001. For subject-specific adaptation, the pre-trained model was fine-tuned on two binary classification tasks as in the first approach. During fine-tuning, the first convolutional layer was frozen to retain the general features learned during pre-training, while the final layer was replaced with a novel fully connected (FC) output layer. Representative training dynamics, including learning curves and fold-specific cross-validation performance for a typical subject, are detailed in Supplementary Fig. S3.

In summary, we constructed a total of five classifiers for each subject: four EEG-based classifiers (two tasks *×* two approaches) and one pose-based classifier. All training and post-analysis were conducted on a workstation with the following specifications: Intel Core i7 CPU, 128 GB RAM, Quadro P6000 GPUs, running Ubuntu 16.04 LTS. All models were implemented using PyTorch v1.12.1^74^.

### Cross-validation

To avoid data leakage, model evaluation was performed using subject-wise cross-validation with predefined folds (8 folds in total). In each fold, data were split into training and test sets, and test data were never used during model selection. Hyperparameters were tuned exclusively within the training data, which were further divided into training and validation subsets (9:1 ratio). The same folds were consistently reused across pose-based, EEG-based, and ensemble classifiers to ensure a fair comparison. Final performance metrics were computed by aggregating results across folds.

To mitigate class imbalance, model training used class-weighted loss functions together with oversampling applied only to the training data within each fold. Class weighting adjusts the loss contribution of minority classes at the optimization level, while oversampling increases the effective representation of minority-class samples during mini-batch training. Their combined use is common in small, imbalanced datasets and was adopted here to stabilize training and reduce sensitivity to subject-specific class imbalance. Oversampling was performed after the train–test split, ensuring that no duplicated samples appeared in validation or test sets, thereby avoiding optimistic bias. We verified that model performance was not driven by duplicated samples and was consistent across folds, even without strict stratification.

### Classifier evaluation

The evaluation criterion is the macro-averaged F1 score. Given the imbalanced nature of our dataset, accuracy can be misleading as it tends to be biased toward the majority class. The F1 score, being the harmonic mean of precision and recall, provides a more balanced and representative measure of our model’s performance on both classes, making it the most suitable metric for our study. The macro-averaged F1 score is given by the following equation:

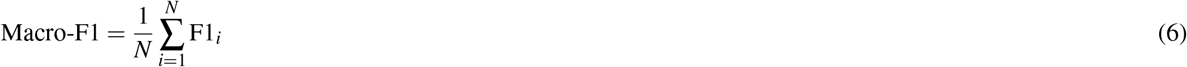

where *N* is the number of classes, F1_*i*_ is the F1 score for the *i*^*th*^ class. For reference, we have included the precision, recall, and accuracy metrics for the final ensemble model, the EEG-based classifier, and the pose-based classifier in Supplementary Tables S3–5, respectively. These measures are given as follows:

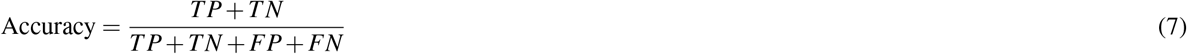

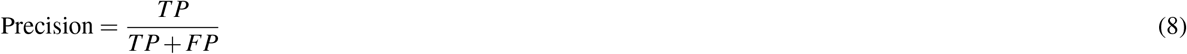

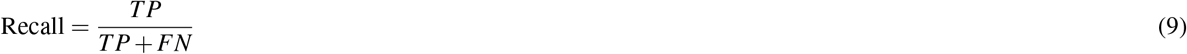

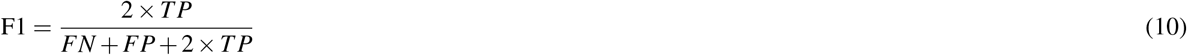

where TP denotes the true positives (correctly identifying the presence of an attack), TN denotes true negatives (correctly identifying the absence of an attack), FN is false negatives (incorrectly identifying the absence of an attack), and FP is false positives (incorrectly identifying the presence of an attack).

To quantify uncertainty given the limited sample size, 95% CIs were estimated using nonparametric bootstrapping across subjects (10,000 resamples). Confidence intervals were computed for macro F1 scores and ensemble gains relative to EEG-only and pose-only classifiers.

Confusion matrices were also computed independently for each subject, normalized row-wise to account for class imbalance, and then averaged across subjects to visualize modality-specific and ensemble error patterns. Per-subject confusion matrices are provided in Supplementary Fig. S4.

### Ensemble approach

We evaluated four ensemble strategies: ensembling with a Decision Tree, stacking with Logistic Regression, blending with a K-Nearest Neighbor (KNN) classifier, and ensembling with Gradient Boosting. Methodologically, the use of Logistic Regression as the final meta-model serves as a direct implementation of the stacking technique, whereas the KNN classifier, when weighted by the inverse distance, acts as a locally weighted voting scheme. We used the Optuna library^75^ to search for the parameters specified below; all other parameters were kept at their default values provided by the scikit-learn library.

- **Decision Tree:** the following parameters were optimized: criterion (“entropy” or “logistic loss”), max depth (from 3 to 20), and splitter strategy (“best” or “random”).
- **Logistic Regression:** we used the Newton-Cholesky solver which is suitable when *n*_*samples*_ *>> n* _*f eatures*_. The maximum number of iterations was set to 10,000, with a tolerance of 10^−7^.The regularization strength (C value) was tuned across values ranging from 10^−8^ to 10^3^ (logarithmic scale).
- **KNN:** Points were weighted by the inverse of their distance to the query point. The number of neighbors was searched over the set: [2, 3, 4, 5, 10, 15, 20, 25, 30, 40, 50, 60, 80, 100, 110].
- **Gradient Boosting:** the learning rate was tuned across values ranging from 10^−6^ to 10 (logarithmic scale). The maximum tree depth and the number of estimators were each varied from 2 to 5.

All strategies were assessed using 8-fold cross-validation. To ensure ensemble quality, we implement a two-stage model selection pipeline. First, we excluded the models that have the macro-averaged F1 score below the chance level (0.5). Second, for each fold, we applied Chi-squared feature selection method on the training set , iterating the number of components (*k*) from 2 to the total number of available models. For each value of k value, we performed hyperparameter tuning, using an internal 10-fold cross-validation split of the training set, to identify the optimal ensemble configuration and its corresponding F1 score. Finally, the combination of k and the best-performing ensemble strategy was used to train the model on the full training set of the respective fold.

### Statistical Analysis

Statistical analyses were conducted in Python v3.10 using the NumPy, SciPy, Pymer4, and scikit-learn libraries. A logistic mixed-effects model was implemented to evaluate the overall effect of the ensemble procedure on decoding performance while accounting for inter-subject variability. For trial *i* belonging to participant *j*, the model is formulated as:

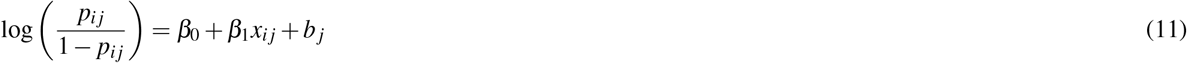

where *p*_*i j*_ = *P*(*Y*_*i j*_ = 1) represents the probability of a correct prediction, *x*_*i j*_ is a binary indicator variable for the model type (1 for ensemble, 0 for baseline), *β*_0_ is the fixed intercept, *β*_1_ is the fixed-effect coefficient for the model condition, and 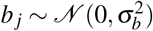 represents the random intercept for participant *j* ∈ {1, … , 9}.

To explore individual differences, we evaluated Spearman’s rank correlation (*ρ*) between each participant’s ensemble performance (macro-F1) and self-reported experience level. Spearman’s correlation was selected due to the ordinal nature of the survey metrics (encoded as ranks 1–4, from lowest to highest) without assuming a linear relationship with decoding accuracy.

## Results

### Behavioral analysis

First, we examine the behavioral tendencies of nine players selected from the public Real World Table Tennis dataset^24^. The type of attack was determined based on the landing region of the table tennis ball on the opponent’s side of the table (Fig. 2a), which was divided into clearly distinguishable left and right zones. These regions were defined such that an observer can unambiguously classify the direction of the ball’s trajectory. Interestingly, most players exhibited a tendency to attack toward the left side (Fig. 2b), with fewer attacks directed toward the right. Although the original dataset documentation did not explicitly detail how handedness influenced this bias, it should be noted that all selected participants in this study are right-handed. “In-between” attack had the highest count; however, these were excluded from our main analysis due to the ambiguity in determining the true attack direction. This observation highlights the importance of carefully considering both model design and evaluation metrics, as the dataset is inherently imbalanced.

**Figure 2.**
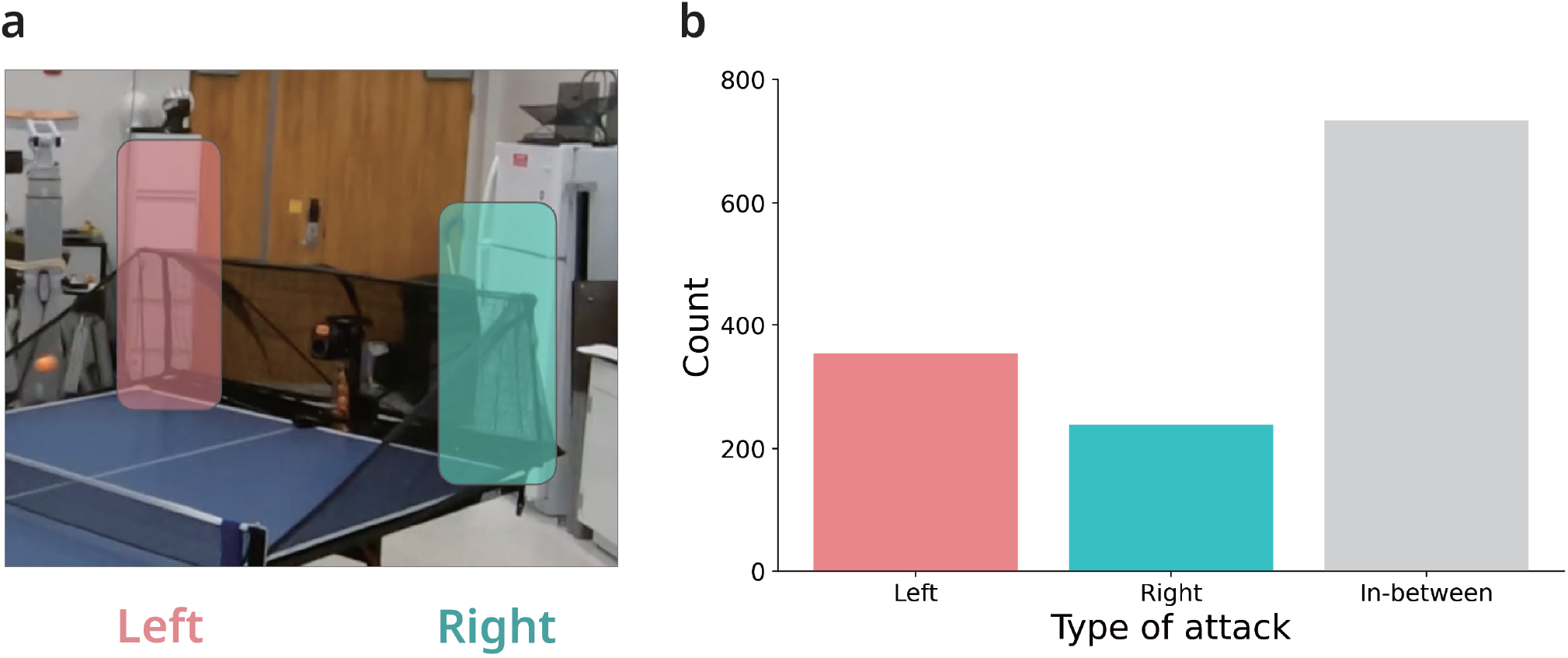
(a) Definition of the “left” and “right” zones. (b) Average count of attack types (left, right, and in-between) aggregated across all subjects.

### Pose-based classifier

To create the pose-based classifier, we employed a simple SGD classifier. Player poses were extracted using the OpenPose model^57^ (Fig. 3a). However, we found that it is visually impossible to recognize the player’s intention at this level of image detail. The original outcomes of the OpenPose model lack information about the spatial relationships between body parts— factors that we believe are crucial for action recognition. To address this limitation, we proposed additional features such as link positions (*x*_*l*_, *y*_*l*_), orientations (*θ*), and lengths (*L*) (Fig. 3b; See Methods for more details). Our preliminary results indicated that while the traditional model struggled to achieve macro-averaged F1 scores above 0.5, the proposed model consistently exceeded this threshold across all participants (averaged F1 score increases by 2.3%), statistically above the chance level (Fig. 3c; one-sample *t*-test, *p* = 0.029). Although the difference between two models did not reach conventional significance (*p* = 0.052, paired *t*-test), the functional improvement highlights the effectiveness of the proposed feature extraction pipeline. Because the timing of pose capture plays a critical role in intention decoding, we determined the appropriate timing by performing time-course analysis with pose extracted at six different time points prior to ball impact: 0 s, −100 ms, −200 ms, −500 ms, −1 s, and − 2 s. The optimal timing was found to be − 100 ms before impact (average macro-F1 score = 0.536 across all subjects; Fig. 4a). Notably, classification performance dropped below chance level when the pose data were taken at more than 1 s prior to impact. Figure 4b shows that 7 out of 9 individual predictors performed well in the left-out test set.

**Figure 3.**
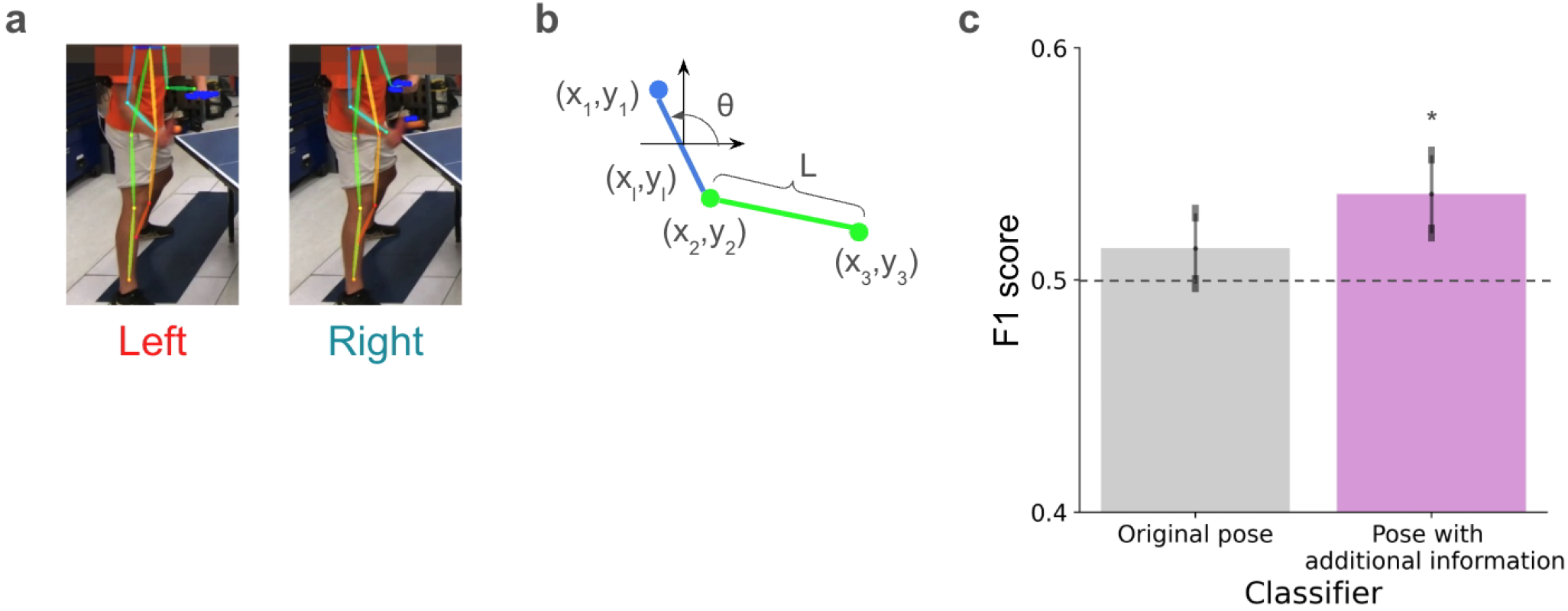
(a) Pose extracted by the OpenPose model during left and right attacks, overlaid on the original video frame. (b) Additional pose features: link positions (*x*_*l*_, *y*_*l*_), orientations (*θ*), and lengths (*L*). (c) Comparison of classification performance between the model using only the original OpenPose outputs (left) and the model incorporating additional features (right). Performance of the proposed model (right) was significantly above chance-level value (one-sample *t*-test, *∗p* < 0.05). The dashed line indicates chance-level performance. Error bars represent the standard error of the mean (SEM).

**Figure 4.**
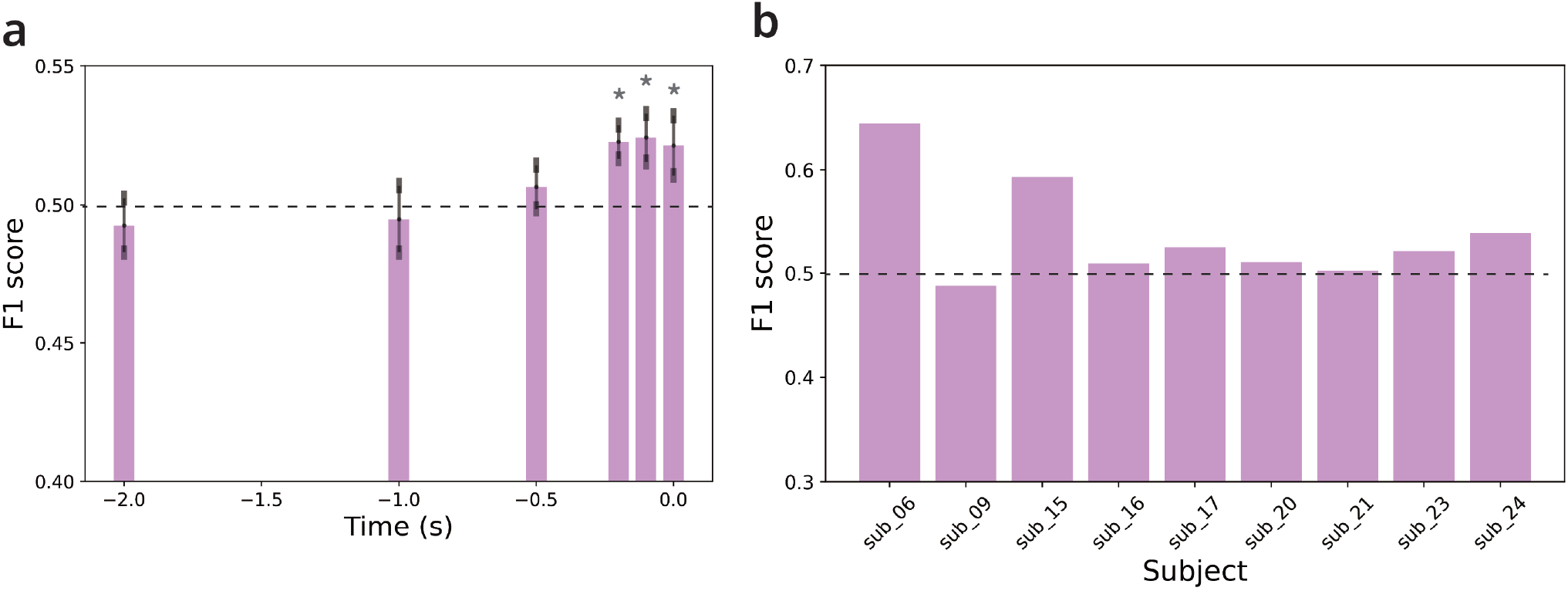
(a) Time-course analysis for the pose-based classifier. The dashed line indicates chance-level performance. \**p* < 0.05; one-sample *t*-test comparing performance against the chance level. Error bars represent the SEM. (b) Test performance of all subject-specific classifiers at −100 ms before impact, the identified optimal timing. Subject indices correspond to their original identifiers in the dataset.

### EEG-based classifier

Consistent with prior motor preparation studies, a moderate *µ*/*β* desynchronization was observed over sensorimotor electrodes C3 and C4 preceding impact (Supplementary Fig. S1). Both frequency bands exhibited a shared temporal profile across hemispheres, characterized by an early power increase peaking around −1.5 s followed by a pronounced desynchronization (ERD) immediately approaching the point of impact (*t* = 0 s). This robust modulation confirms that the recorded EEG signals capture physiologically plausible motor-preparatory dynamics. Crucially, no dissociation in peak timing and amplitude between left and right attacks was observed at the group level. The absence of overt single-channel differences underscores that the intention-decoding task is highly non-trivial, implying that the downstream classifiers must leverage subtle, distributed multivariate patterns across multiple electrode combinations to achieve successful classification.

To create the EEG-based classifier, we adopted the EEGNet architecture (originally supports 64 channels of EEG data)^13^ (Fig. 5a), modified to accommodate all 120 EEG channels in our dataset. To address the class imbalance, we incorporated precomputed class weights into the loss function together with oversampling applied only to the training data within each fold. These class weights were calculated individually for each subject, as their behaviors were not consistent across the dataset (see Methods for more details).

**Figure 5.**
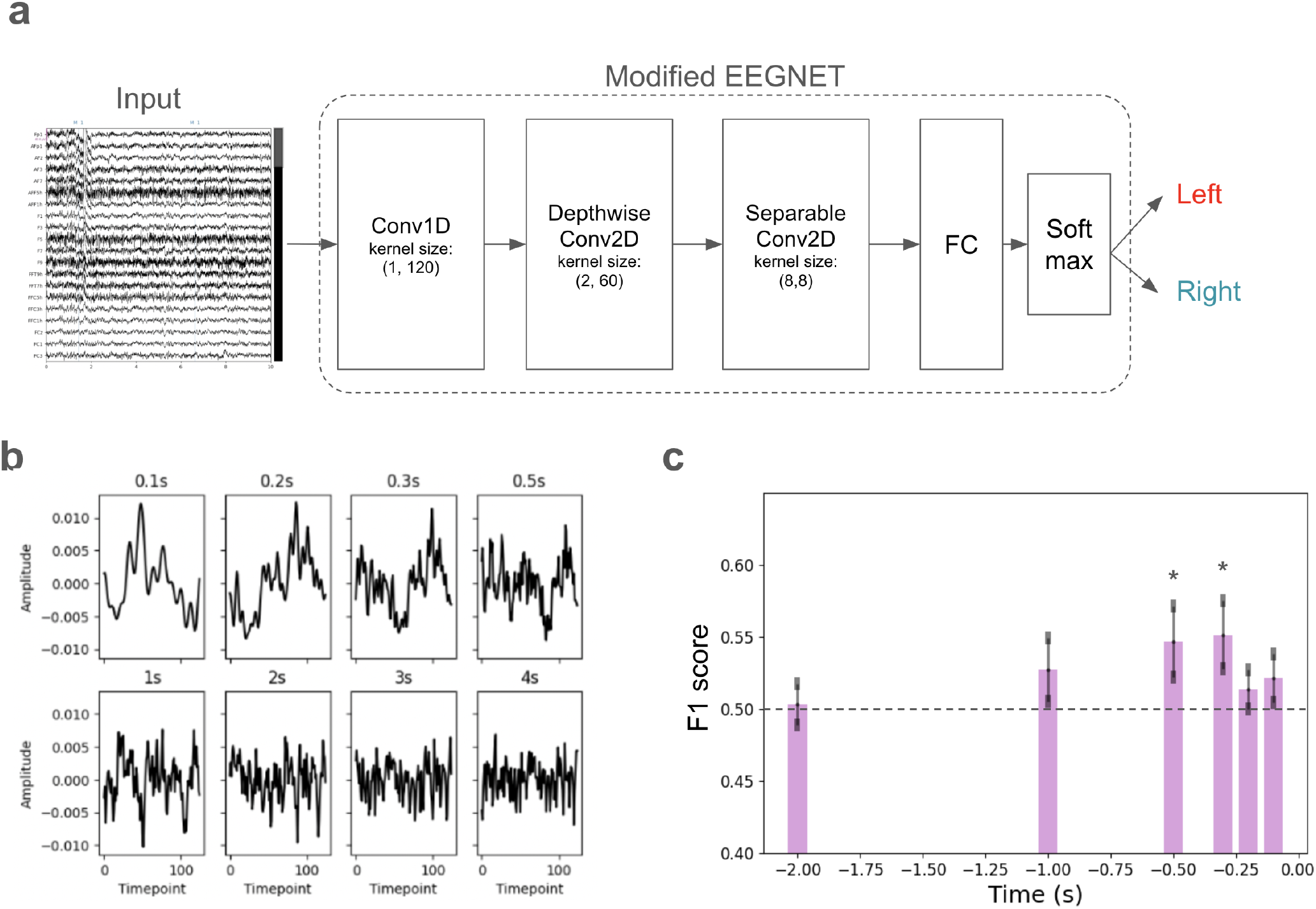
(a) Architecture of our EEG-based classifier, adapted from EEGNET^13^. (b) Examples of EEG signals at various time points prior to impact. The titles of subplots indicate the timing relative to impact. (c) Classification performance at each time point. The dashed line indicates chance-level performance. Statistical comparisons show that the performance was significantly above the chance-level value at −300 ms and − 500 ms before impact (one-sample *t*-test, *∗ p* < 0.05). Error bars represent the SEM.

Following a similar approach to pose-based analysis, we evaluated multiple time points to identify the optimal EEG window for intention decoding. EEG segments were extracted starting at −100 ms, −200 ms, −300 ms, −500 ms, − 1 s, −2 s before impact (Fig. 5b), with − 3 s and −4 s windows included to demonstrate the complexity of longer intervals. The windows end exactly at the impact marker. Here we found that the best performance was achieved at −500 ms before impact (average F1 score = 0.542; Fig. 5c). F1 scores at both −300 and −500 ms before impact were significantly above the chance level (one-sample *t*-test; *t*(8) = 2.82; *p* = 0.011), while other timings did not. These results suggest that the EEG-based classifier can decode intention earlier than pose-based model and possibly ball-tracking model.

As an initial validation to motivate our choice of architecture, we compared the modified EEGNet with a conventional SGD classifier using ElasticNet regularization, as well as with recently proposed Transformer-based models, including EEGConformer^19^ and CTNet^20^ (Fig. 6a). We found that all neural network-based models outperformed the SGD classifier with ElasticNet regularization. Further analysis confirmed the superior performance of the modified EEGNet over a conventional SGD classifier with ElasticNet regularization (Supplementary Fig. S5; paired *t*-test, *t*(8) = 2.09, *p* = 0.034). Among the neural networks, EEGNet achieved slightly better performance. We believe this outcome is largely attributable to the sample-size limitations in the current dataset (Supplementary Table S2). The Transformer-based models, with their substantially larger number of parameters (Fig. 6b), require more training data to fully realize their advantages and may therefore be more susceptible to overfitting under the present conditions.

**Figure 6.**
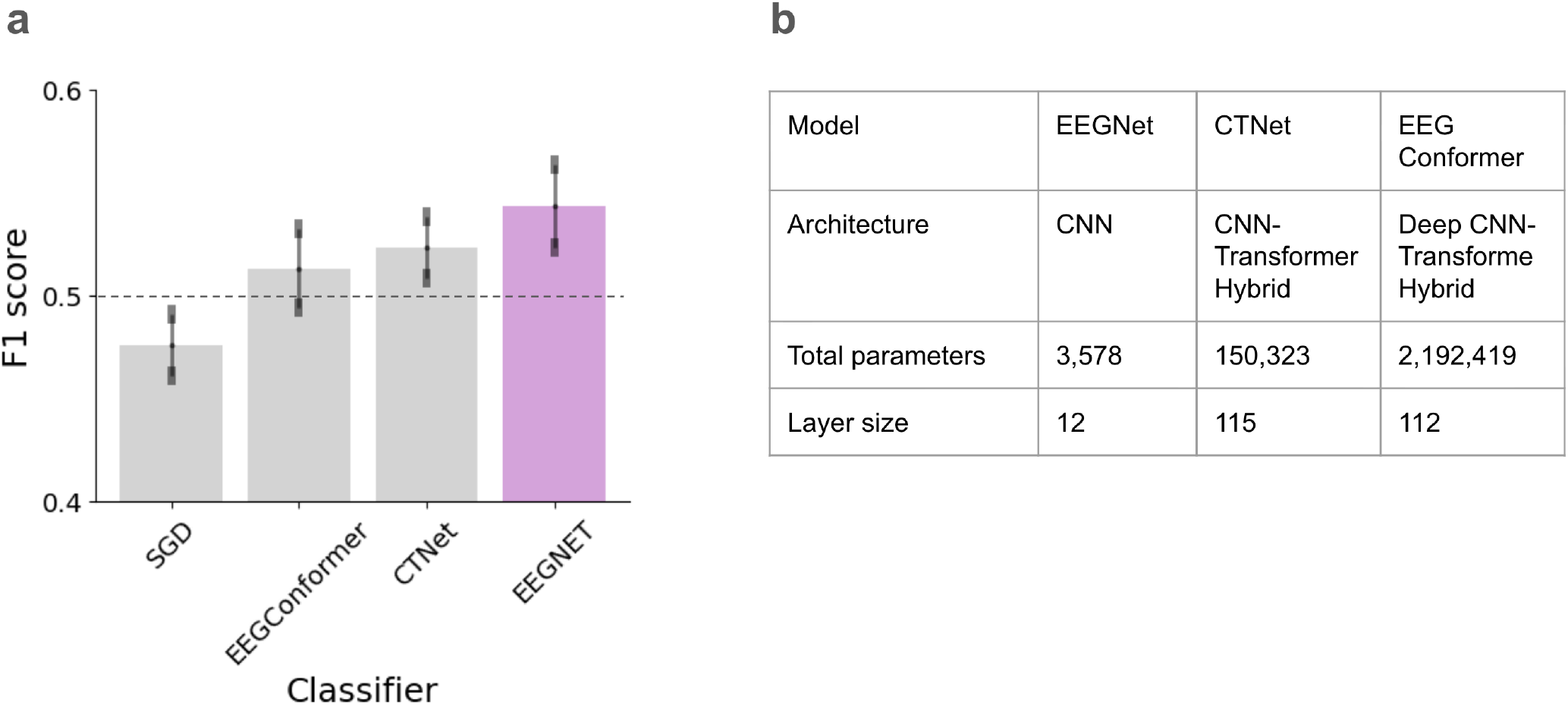
(a) Comparisons of performance between EEGNet and recently proposed models (EEGConformer, CTNet) at −500 ms before impact. The dashed line indicates chance-level performance. Error bars represent the SEM. (b) Summary of the evaluated network architectures.

Having established EEGNet as an appropriate backbone, we next describe the detailed implementation of the EEG-based classifiers. Given the strong class imbalance in the dataset, conventional training with a large neural network architecture would likely result in a model biased toward the major class (i.e., “left”). To avoid this, we trained two separate binary classifiers: one for “Left vs. Others” (LvsO) and another for “Right vs. Others” (RvsO) (Fig. 7a). We explored two distinct approaches to create the EEG classifier, i.e., direct training approach and transfer learning approach (Fig. 7b). To construct the group model, we adopted a leave-one-subject-out (LOSO) scheme, in which data from all subjects except the target were used for pre-training. The resulting model was then fine-tuned using the left-out subject’s data.

**Figure 7.**
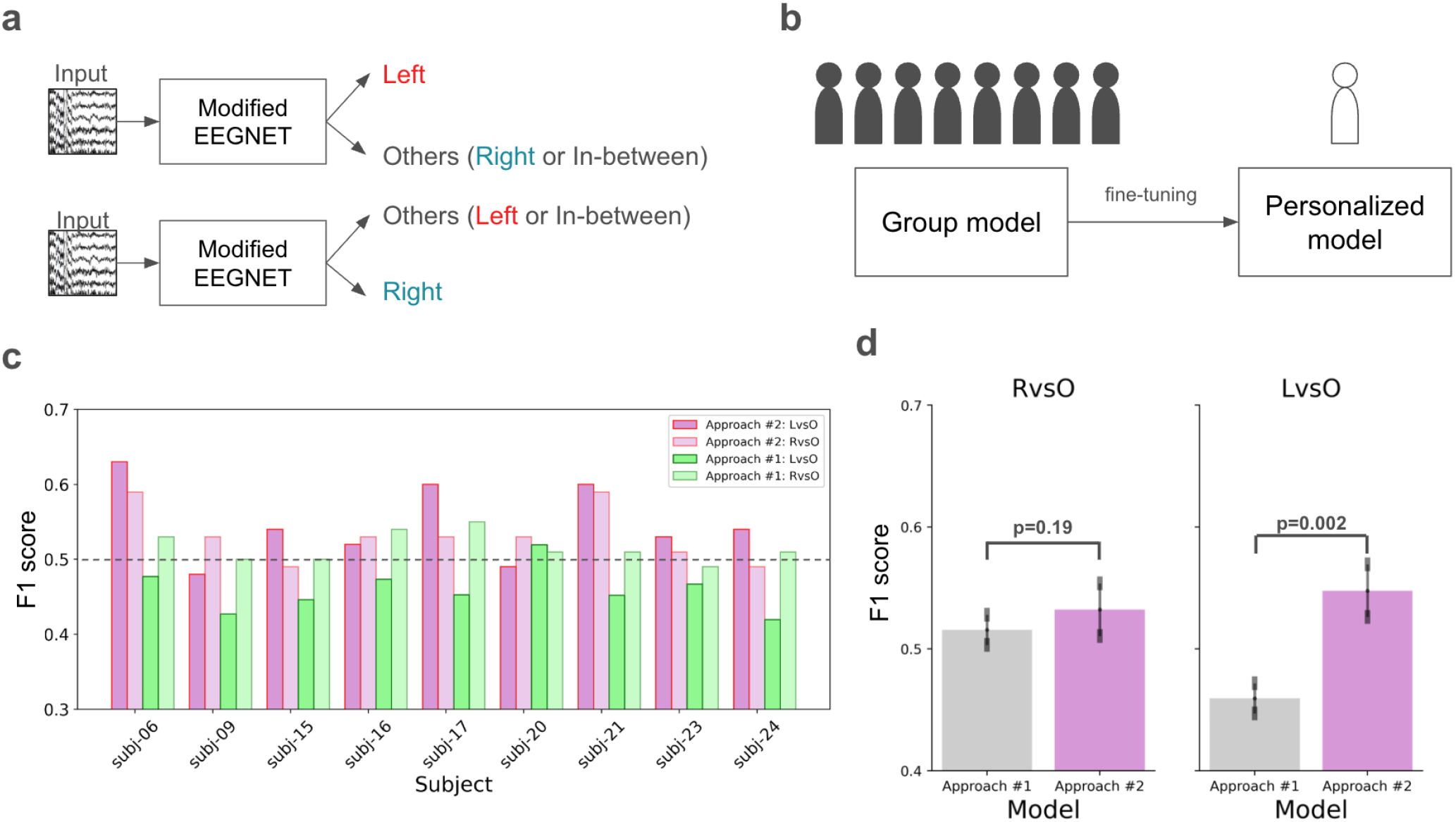
(a) Schematic of the first approach: training two separate models (“Left vs. Others” and “Right vs. Others”). (b) Schematic of the second approach using transfer learning from a group-level model. (c) Performance summary of all EEG-based models; the dashed line indicates chance-level performance. (d) Statistical comparison between the first and second approaches via paired *t*-test. Error bars represent the SEM.

Figure 7c shows the F1 scores for all models. Overall, the LvsO models consistently outperformed the RvsO models across participants. Notably, models trained using the second approach (transfer learning) significantly outperformed those trained using the first approach (subject-specific training) for the LvsO classification task (paired *t*-test; *t*(8) = 4.35, *p* = 0.002; Fig. 7d). The improvement was substantial, increasing the mean F1 score by 8.8% in the LvsO task. In contrast, no significant difference was observed between the two approaches for the RvsO task (paired *t*-test; *t*(8) = 1.40, *p* = 0.19), although the mean F1 score still saw a modest 1.6% increase. Taken together, these results suggest that the transfer learning approach yields superior performance compared to the direct training approach, confirming our hypothesis.

### Intention decoding with ensemble approach

Subsequently, we evaluated the effectiveness of the ensemble that combines pose-based and EEG-based classifiers. Four algorithms–Logistic Regression, KNN, Gradient Boosting, and Decision Tree algorithms–were compared, and the optimal one was chosen by cross-validation (Fig. 8a). Within each fold, we further performed model-selection based on their Chi-squared score computed on the training set.

**Figure 8.**
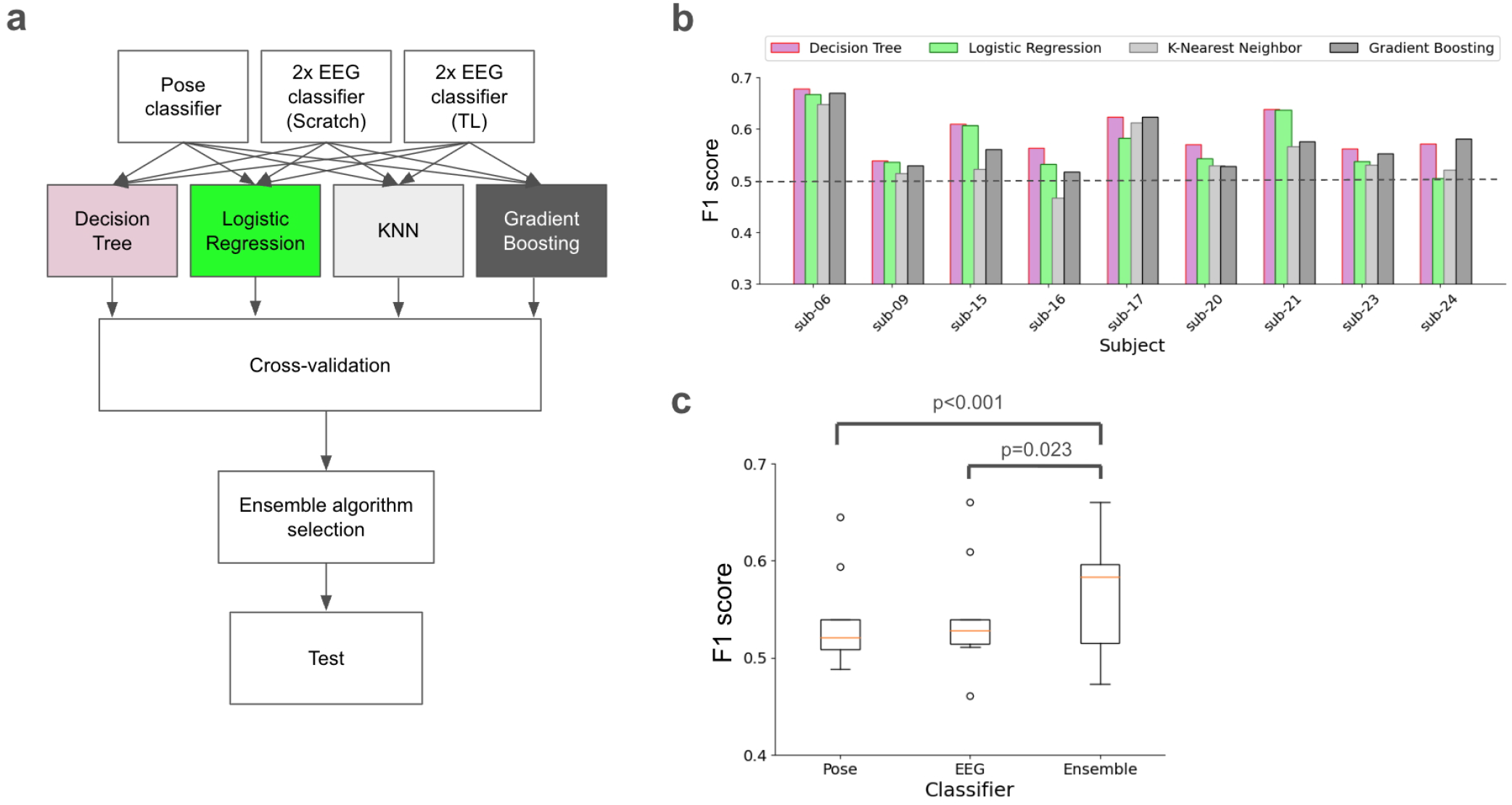
(a) Schematic overview of the ensemble approach. (b) Summary of the employed strategies and corresponding macro F1 scores for each subject on the hold-out test set; the dashed line indicates chance-level performance. (c) Performance comparison of the ensemble classifier against its associated pose- and EEG-based classifiers (statistically evaluated via a logistic mixed-effects model). Orange lines indicate median values.

Macro-F1 scores for all ensemble techniques are shown in Fig. 8b; the Decision Tree algorithm delivered the best performance across most scenarios. The ensemble model achieved an average macro-F1 score of 0.563, with a bootstrapped 95% CI of [0.523, 0.603], reflecting notable inter-subject variability (range across subjects: 0.47–0.65). Compared to single-modality baselines, the ensemble demonstrated a consistent performance advantage (Fig. 8c). Specifically, the ensemble improved over the pose-only classifier by Δ*F*1 = +0.026, with a 95% bootstrapped CI of [0.00, 0.057], supported by a significant logistic mixed-effects model (*z* = 4.48, *p* < 0.001). The improvement over the EEG-only classifier was smaller (Δ*F*1 = +0.02) and exhibited greater uncertainty (95% CI [−0.016, 0.062]), though it remained significant under the mixed-effects analysis (*z* = 2.269, *p* = 0.023), suggesting a modest but variable ensemble benefit across subjects. This divergence highlights that while the ensemble benefit is variable at the individual subject level due to the limited sample size (*N* = 9), a significant positive trend emerges when accounting for nested trial-level dynamics and inter-participant variance simultaneously. The median F1 score increased by 0.06 from the EEG-only classifier and 0.062 from the pose-only classifier. No significant differences were observed between pose-only and EEG-only classifiers (*z* = 1.827, *p* = 0.068). Taken together, these findings suggest that the multi-modal ensemble strategy provides a valuable advantage for intention decoding.

Figure 9 presents the group-level normalized confusion matrices for the pose-based, EEG-based, and ensemble classifiers. The pose-based classifier showed higher accuracy for right attacks (53.97%) than for left attacks, whereas the EEG-based classifier exhibited the opposite pattern, with improved performance for left attacks (61.97%) but reduced accuracy for right-class trials (47.54%). The ensemble classifier reduced misclassification rates for both classes, yielding the most balanced prediction profile across left and right intentions. This reduction in class-specific errors is consistent with the observed improvements in macro-F1 score for the ensemble model (Δ*F*1 = +0.03 over pose-only and Δ*F*1 = +0.02 over EEG-only; Fig. 8c), reflecting improved overall precision–recall trade-offs across classes. Per-subject confusion matrices are provided in Supplementary Fig. S4.

**Figure 9.**
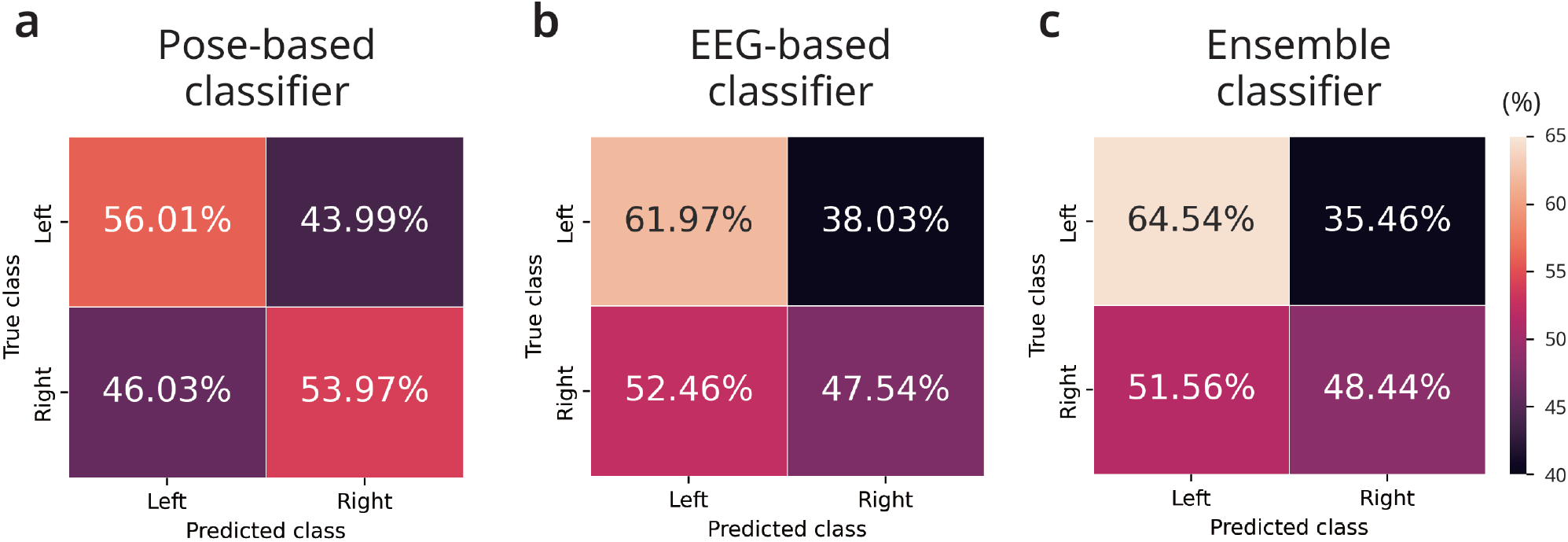
Group-level confusion matrices (in %) for the (a) pose-based classifier, (b) EEG-based classifier, and (c) ensemble classifier. Values represent mean classification percentages across subjects, with rows normalized by the true class (Left vs. Right).

To examine whether inter-individual differences in ensemble performance were related to player experience, we computed Spearman’s rank correlations between survey responses and per-subject macro-F1 scores. We observed a moderate positive association between self-reported playing frequency (Q1) and ensemble model performance (Spearman’s *ρ* = 0.46); however, this effect did not reach statistical significance (*p* = 0.21), likely due to the limited number of participants. No association was found between tournament history (Q2) and model performance (*ρ* = −0.10, *p* = 0.79). A stronger negative correlation was observed between self-level assessment (Q3) and model performance (*ρ* = −0.55, *p* = 0.12); however, this effect was driven by a single influential outlier rather than a consistent overall trend (see Supplementary Fig. S6). Overall, these results suggest possible inter-individual trends that warrant further investigation in larger cohorts.

## Discussion

In this study, we proposed a novel intention decoding approach that uses both pose-based and EEG-based classifiers. Our results demonstrated the superiority of the ensemble model over its individual components, suggesting a potential improvement over the performance reported by pure pose-based classifiers in previous studies^34,35^. The complementary error patterns observed in Fig. 9 suggest that pose and EEG signals capture partially non-overlapping information, which likely contributes to the ensemble’s improved macro-F1 score. It should be noted that our approach was successfully applied to a real-world motor task, predicting intention several hundred milliseconds before impact (speficically, − 500 ms for EEG and −100 ms for pose). This is a crucial departure from conventional motor intention decoding studies, which typically rely on the motor imagery paradigm^51,63,76–78^. Compared to our real-world tasks, motor imagery is methodologically simpler but less consistent with how the brain actually plans motion.

A major advantage of the ensemble framework is its modularity, enabling the independent incorporation of diverse data modalities. This scalability allows for future extensions to include other physiological signals, such as information from the IMUs or electromyography (EMG). This approach aligns with the growing trends of multi-modal ensemble methods across various domains, including biometric recognition^46^, human activity recognition^79^, Parkinson quantification^47^, and clinical readmission^48^. It should be noted that individual differences, originating from players’ experience, playing styles, or decision-making policy, can influence decoding performance. This explains the variation in optimal ensemble strategies across subjects shown in Fig. 8b. For a real-world application, one should always consider fine-tuning the ensemble with an individual-specific dataset for better decoding accuracy. While our approach adopts the decision-level ensemble, feature-level fusion approaches have been explored in other domains^66,67^. Although their application to movement-intensive EEG settings remains challenging due to data scale and noise sensitivity, the feature-level fusion could be leveraged as an independent module to improve the system performance in future work.

Interpreting participants’ intentions is a highly challenging task. Even the participants themselves may struggle to pinpoint the precise moment when their intentions emerge, particularly at the sub-second level. Additionally, there is a moderate likelihood of an “intention-behavior” gap^80,81^, where a participant’s actions may not match their underlying intentions. To date, no solution fully addresses this issue. One potential approach is to predict the success of participant intention based on their facial responses. Another approach, while carrying a potential risk of distraction, would be to ask participants to verbally describe their intention after the ball is served and record it via a microphone. Alternatively, one could design an experiment where participants preregister their intentions (e.g., left only, right only, or left-right sequentially) in advance. This would ensure a well-balanced dataset, thereby improving the precision of the classifiers. However, these approaches were not feasible in our study due to the masking of facial features for privacy protection and the lack of voice recordings. Despite these limitations, our model still achieved robust performance, underscoring the benefit of integrating both pose and EEG data.

Previous studies have shown that pose information can be useful for decoding intentions from movement kinematics^53^ and for strategic analysis^34^. In this study, we demonstrated that pose information can act as an early indicator of left-right attack intentions. With the inclusion of additional kinematic features (i.e., joint orientations and limb lengths), the pose-based classifier achieved effective performance, even though the task remains visually challenging for a human observer given the same instantaneous information (Fig. 4). Here, we focused on static pose features to establish a stable and interpretable baseline that is less sensitive to frame-rate variability and pose estimation noise, which can be pronounced during high-speed movements in table tennis. Incorporating higher-order temporal derivatives, such as velocities and accelerations, is a promising direction and may further enhance predictive power in future work.

Furthermore, we found that EEG signals could predict left-right attacks as early as −500 ms before impact. This aligns with our expectations, given that brain signals naturally precede physical actions. However, the performance of EEG-based classifiers was not significantly higher than that of the pose-based classifiers, despite requiring more training effort. A possible explanation for this could be the presence of intrinsic artifacts in the EEG data^82^. Although the EEG signals were meticulously preprocessed (see Method and original paper^24^ for details), it is impossible to completely eliminate noise from EEG recordings. Moreover, since the original experiment was not specifically designed for intention decoding, the EEG signals likely contain a mixture of unrelated neural activity, not limited to intention alone.

The application of CNNs for target prediction has recently gained attention^32^, particularly in the domains of image and video analysis. Here, we demonstrated that CNNs can also be effectively applied to intention decoding using raw EEG signals, achieving performance comparable to that of the pose-based classifier. We additionally tested many variants of CNN architectures for EEG classification. A critical requirement for the success of ensemble approaches is that their component models make diverse and uncorrelated errors. However, due to the inherent class imbalance in the dataset, favoring left-side attacks (Fig. 2b), naive training of the models tends to yield similar error patterns. To mitigate this, we introduced a “separate- to-training” strategy for the EEG-based classifiers, which compels the classifiers to become sensitive to either left or right attack types, thereby promoting diversity in decision boundaries and improving the overall robustness of the final ensemble.

Furthermore, we also investigated the impact of training methodology on model performance by comparing direct training and transfer-learning approaches. Our results indicate that transfer learning yielded superior performance. This finding may be attributed to the transfer learning approach using a larger-scale dataset to train the base models, which improves the initialization of shallow layers and enhances the generalization capabilities when fine-tuning. This pre-training step provides a more robust foundation for learning than training on individual, limited datasets. Future work could explore extending CNN applications to pose analysis. For instance, modifying the OpenPose model to directly predict the left-right attack intentions or unifying it with the two EEG classification approaches explored in this study could be valuable. Such a combined model could potentially leverage the complex interaction between kinematic and neural information for a more comprehensive and accurate prediction of attack intention.

We emphasize that ERD/ERS analysis is presented only as a qualitative sanity check. The primary contribution of this work lies in predictive decoding rather than neurophysiological modeling. Therefore, a more detailed causal or source-level analysis is beyond the scope of the present study and should be considered in future work. Our findings indicated that intentions become most detectable approximately −100 ms before ball impact via pose data, and approximately −500 ms before impact via EEG signals. While the classification performance decreased beyond these windows, we argue that the actual intention underlying an action likely emerges earlier and is linked to broader tactical and strategic considerations. This aligns with the concept of multiple levels of intention, as theorists distinguish “intention” into “immediate/proximal intentions” and “distal intentions” based on the temporal distance of the planned action^83^. The intentions decoded in this study are likely “proximal”; further investigation will be necessary to explore the characteristics and detectability of the “distal intentions”.

From an applied perspective, our proposed method holds promise for advancing neural prosthetic systems^2^ and can be deployed in sports training as a neurofeedback tool to help the player improve their skills. However, we acknowledge that the current implementation is not designed for real-time deployment. In particular, the use of high-density EEG (120 channels), offline preprocessing pipelines, and computationally intensive pose estimation introduces substantial latency that limits immediate real-time feasibility. To mitigate these practical limitations, we recommend that future work implements the channel selection/analysis techniques, such as the Sparse Common Spatial Pattern algorithm^84^, or Grad-CAM^63^, to decrease both acquisition burden on participant’s head and computational cost. On the kinematic side, replacing the current pose extraction pipeline with lightweight and faster pose estimators^85^, together with edge or on-device EEG processing strategies^15^, could further reduce end-to-end latency. While these adaptations were beyond the scope of the present study, they represent important directions for future work aimed at enabling real-time or near–real-time applications under movement-intensive conditions.

### Limitations

Our simplified binary classification of left and right attacks captures only a limited aspect of the complex decision-making in a table tennis game. Although the proposed framework demonstrates performance significantly above chance level, its current form offers limited immediate practical utility. The macro-F1 score is highly varied across subjects, with a maximum of 0.65 and a minimum of 0.473. This variance suggests substantial inter-individual differences, potentially arising from differences in playing style, movement consistency, or neurophysiological signal quality, which may limit the generalization to unseen subjects. We additionally explored whether inter-individual differences in decoding performance were related to self-reported playing experience. Although moderate correlations were observed for some survey measures, none reached statistical significance, likely due to the limited number of participants. Future work with larger cohorts will be needed to examine the relationship between decoding performance and player expertise or behavioral factors.

The number of samples per subject is also small (ranging from 464 to 769 samples) and highly imbalanced, thus limiting the model’s generalization. To enhance model precision and practical applicability, we suggest that future work focuses on the collection of large-scale data with more detailed and nuanced labels. Ideally, the dataset should be collected from individuals who are intensively trained in a professional environment. Additionally, incorporating richer pose information (e.g., labels for backhand flip action, chop action, etc.^34^) could be valuable for future exploration, as these actions convey critical details about the player’s intentions and technique.

Another limitation lies in the labeling process of the dataset. The current annotations were manually provided by an external observer, assuming that the participants executed their intended action flawlessly. Although care was taken to ensure label accuracy, allowing participants to self-report their intentions would yield a more reliable ground truth and improve the model’s interpretability.

Furthermore, the present study relied exclusively on pre-recorded and preprocessed data; critical engineering challenges regarding real-time implementation, such as processing cycle time, strict cross-modal synchronization, and real-time artifact removal, remain unaddressed. More effort is required to advance this research to the next phase, i.e., real-time, closed-loop intention decoding^86^, in future work.

Finally, we did not explicitly remove or model motion-related covariates when analyzing EEG signals recorded during active gameplay. Although artifact removal procedures provided with the dataset reduce muscle and motion contamination, residual movement effects may still contribute to decoding performance. While comparisons between EEG-only, pose-only, and ensemble models suggest that EEG retains predictive information beyond pose features, this does not constitute causal disentanglement. Future studies should apply explicit motion-covariate regression or controlled experimental designs to more rigorously separate neural intention signals from movement-related artifacts.

## Conclusion

In this study, we proposed a multi-modal ensemble framework for intention decoding using EEG signals and kinematic (pose) information. Using a public real-world table tennis dataset, we showed that both modality-specific classifiers can predict left-right attack intentions from time windows up to several hundred milliseconds (−100 ms to −500 ms depending on modality) prior to racket-ball impact. The proposed decision-level ensemble consistently outperformed the individual EEG- and pose-based classifiers, indicating the complementary nature of neural and kinematic signals in movement-intensive scenarios. Although the present evaluation was conducted using offline analysis, the results suggest the potential relevance of this framework for applications such as sports-related neurofeedback and neural prosthetic systems under realistic conditions.

Future work will focus on methodological and engineering optimizations, including sensor reduction, computational efficiency, and low-latency processing, to enable real-time or near-real-time deployment of the proposed approach.

## Data availability

The raw datasets used in the current study are publicly available in the OpenNeuro repository under the following DOI: 10.18112/openneuro.ds004505.v1.0.4. All other materials, generated data, and codes are available from the corresponding author upon reasonable request.

## Acknowledgements (not compulsory)

We thank Dr. Nanami Taketomi and Dr. Yasushi Orihashi from the Clinical Research Center in Hiroshima, Hiroshima University Hospital for their supportive comments in statistical analysis. This work is supported by the following funds: Japan Science and Technology Agency (JST) Moonshot R&D Grant (No. JPMJMS2012) to R.K., Japan Society for the Promotion of Science (JSPS) KAKENHI Grant (No. 24K03243) to J.C, Japan Agency for Medical Research and Development (AMED) Grant (No. JP24gm7010007 and No. JP256f0137011) to J.C., and JSPS KAKENHI Grant (No. 23K17182) to T.Q.P.

## Author contributions statement

Conceptualization: T.Q.P., R.K., J.C. Methodology: T.Q.P., S.S.F, J.C. Investigation: T.Q.P, J.C. Writing - Original Draft: T.Q.P. Writing - Review and Editing: T.Q.P., S.S.F, R.K, J.C. Supervision and Funding Acquisition: R.K., J.C. All authors reviewed the manuscript.

## Competing interests

S.S.F is employed by OneAI, Inc. R.K is the founder and CEO of Araya, Inc., and J.C is employed by Araya, Inc. These affiliations had no involvement in the design, analysis, or interpretation of the results. Other authors declare no competing interests.

## Notes

### Summary of Updates

Reorganized the order of the sections; Corrected the mislabeling of the columns in Table S1; Updated Statistical Analysis section for better clarification

