## Supplemental Materials for "Multi-modal Ensemble Approach for Decoding Player Intentions in Table Tennis"

**Supplementary Information**

### Equivalence of two-class softmax and sigmoid activation

Consider a classifier with two outputs, $z_1$ and $z_2$. The softmax probability for class 1 is:

$$p_1 = \frac{e^{z_1}}{e^{z_1} + e^{z_2}} = \frac{1}{1 + \frac{e^{z_2}}{e^{z_1}}} = \frac{1}{1 + e^{z_2 - z_1}}$$

Let $x = z_1 - z_2$. Then:

$$p_1 = \frac{1}{1 + e^{-x}} = \sigma(x)$$

where $\sigma(x)$ denotes the sigmoid function for a single neuron.

Thus, in the binary case, the softmax output for one class is equivalent to applying a sigmoid function to the logit difference $z_1 - z_2$. Under cross-entropy loss, predictions depend only on this logit difference; therefore, a model with two softmax outputs produces identical decision boundaries to a model with a single sigmoid output neuron. Consequently, both formulations are mathematically equivalent for binary classification.

### Multi-modal Ensemble Approach for Decoding Player Intentions in Table Tennis

| Subject | Age | Gender | Left attacks | Right attacks | In-between attacks | Total |
|---|---|---|---|---|---|---|
| sub-06 | 29 | M | 439 | 265 | 798 | 1502 |
| sub-09 | 22 | M | 389 | 296 | 661 | 1346 |
| sub-15 | 30 | F | 315 | 149 | 938 | 1402 |
| sub-16 | 20 | M | 395 | 233 | 749 | 1377 |
| sub-17 | 18 | M | 448 | 321 | 641 | 1410 |
| sub-20 | 20 | M | 304 | 392 | 685 | 1381 |
| sub-21 | 18 | M | 287 | 214 | 1128 | 1629 |
| sub-23 | 18 | M | 359 | 281 | 731 | 1371 |
| sub-24 | 21 | F | 552 | 131 | 815 | 1498 |

**Table S1.** Participant's demographic data and sample counts per class (Left attacks, right attacks) after the selection process. Left and right attack trials correspond to the primary analysis targets. F: Female, M: Male. Counts for in-between attacks and total samples are reported for reference.

| Subject | Q1 | Q2 | Q3 | Q6 | Q7 | Q9 | EEG | Video | Sample size | Decision |
|---|---|---|---|---|---|---|---|---|---|---|
| sub-01 | a | b | c | d | d | c | Yes | No | N/A | Excluded |
| sub-02 | a | a | b | a | b | a | Yes | No | N/A | Excluded |
| sub-03 | a | a | b | d | d | c | Yes | No | N/A | Excluded |
| sub-04 | a | a | b | d | d | c | Yes | No | N/A | Excluded |
| sub-05 | a | a | b | d | c | c | Yes | No | N/A | Excluded |
| sub-06 | b | a | b | c | b | c | Yes | Yes | 704 | Included |
| sub-07 | a | a | b | a | a | b | Yes | Yes | N/A | Excluded |
| sub-08 | a | a | a | b | b | d | Yes | No | N/A | Excluded |
| sub-09 | a | b | c | d | c | c | Yes | Yes | 685 | Included |
| sub-10 | c | a | b | a | b | b | Yes | Yes | 283 | Excluded (insufficient sample) |
| sub-11 | a | a | b | a | a | b | Yes | Yes | N/A | Excluded |
| sub-12 | a | a | b | a | a | c | Yes | Yes | N/A | Excluded |
| sub-13 | a | a | b | a | b | c | Yes | Yes | N/A | Excluded |
| sub-14 | a | a | b | a | b | b | Yes | Yes | N/A | Excluded |
| sub-15 | c | a | b | a | b | b | Yes | Yes | 464 | Included |
| sub-16 | b | a | b | a | b | b | Yes | Yes | 628 | Included |
| sub-17 | d | a | b | a | b | b | Yes | Yes | 769 | Included |
| sub-18 | b | a | b | a | b | b | Yes | No | N/A | Excluded |
| sub-19 | a | a | b | a | b | b | Yes | Yes | N/A | Excluded |
| sub-20 | b | a | b | d | c | c | Yes | Yes | 696 | Included |
| sub-21 | b | b | b | c | c | b | Yes | Yes | 501 | Included |
| sub-22 | a | a | b | d | c | c | Yes | No | N/A | Excluded |
| sub-23 | b | a | b | a | b | b | Yes | Yes | 640 | Included |
| sub-24 | a | a | b | d | c | c | Yes | Yes | 683 | Included |
| sub-25 | b | a | b | a | b | b | Yes | Yes | 372 | Excluded (insufficient sample) |

**Table S2.** Participant survey responses to key questions, data availability, and the final decision for the analysis. Q1: Playing frequency (Table Tennis). Q2: Tournament history (Table Tennis). Q3: self-level assessment (Table Tennis). Q6: Tournament history (Other Racquet Sports), Q7: Self-assessment (Other Racquet Sports). Ranks for Q1, Q2, Q3, Q6 and Q7 range from "a" to "d", where "d" indicates a higher level of experience or performance. Q9: Primary racquet sport ("a": None, "b": Table Tennis, "c": Tennis, "d": Badminton). N/A indicates not applicable; Sample sizes reflect only left and right attack trials, which constituted the primary analysis targets; in-between attack trials were excluded from the counts. Sample sizes were not counted for participants who did not meet the survey inclusion criteria.

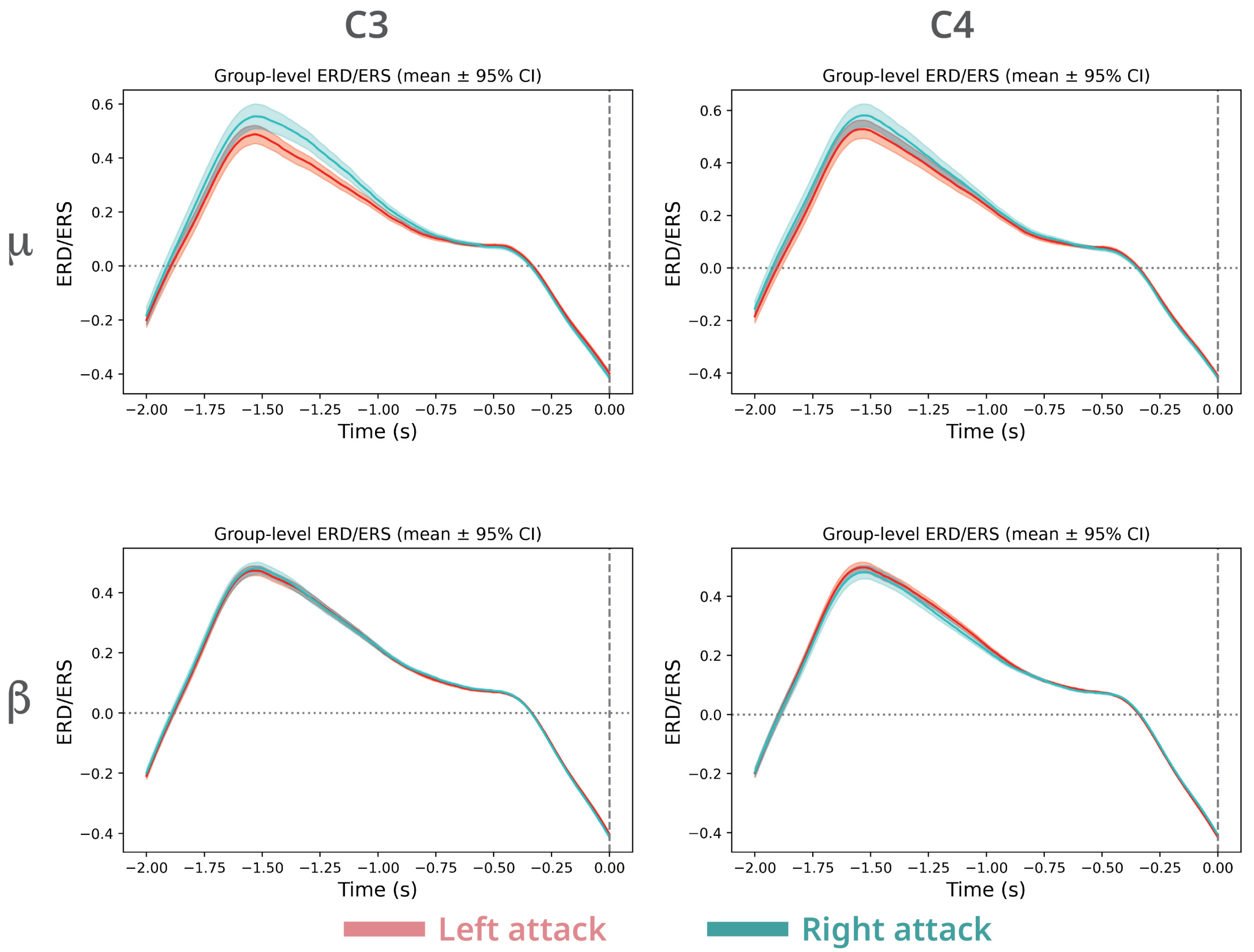


**Figure S1.** Group-averaged event-related desynchronization/synchronization (ERD/ERS) time courses (mean ± 95% confidence interval across subjects) for the $\mu$ (8–13 Hz, top row) and $\beta$ (13–30 Hz, bottom row) frequency bands at electrodes C3 (left) and C4 (right). Time is aligned to racket–ball impact (0 s; vertical dashed line). ERD/ERS values are baseline-normalized, with negative values indicating desynchronization and positive values indicating synchronization. Shaded areas indicate inter-subject uncertainty.

| Layer (type) | Output Shape | Param # |
| --- | --- | --- |
| Conv2d-1 | [-1, 16, 125, 1] | 1,936 |
| ELU-2 | [-1, 16, 125, 1] | 0 |
| BatchNorm2d-3 | [-1, 16, 125, 1] | 32 |
| Dropout-4 | [-1, 16, 125, 1] | 0 |
| ZeroPad2d-5 | [-1, 1, 17, 158] | 0 |
| Conv2d-6 | [-1, 4, 16, 99] | 484 |
| ELU-7 | [-1, 4, 16, 99] | 0 |
| BatchNorm2d-8 | [-1, 4, 16, 99] | 8 |
| Dropout-9 | [-1, 4, 16, 99] | 0 |
| MaxPool2d-10 | [-1, 4, 4, 25] | 0 |
| ZeroPad2d-11 | [-1, 4, 11, 28] | 0 |
| Conv2d-12 | [-1, 4, 4, 21] | 1,028 |
| ELU-13 | [-1, 4, 4, 21] | 0 |
| BatchNorm2d-14 | [-1, 4, 4, 21] | 8 |
| Dropout-15 | [-1, 4, 4, 21] | 0 |
| MaxPool2d-16 | [-1, 4, 2, 5] | 0 |
| Linear-17 | [-1, 2] | 82 |
| Softmax-18 | [-1, 2] | 0 |

**Figure S2.** Architecture of the modified EEGNet model with a dropout probability of 0.25.

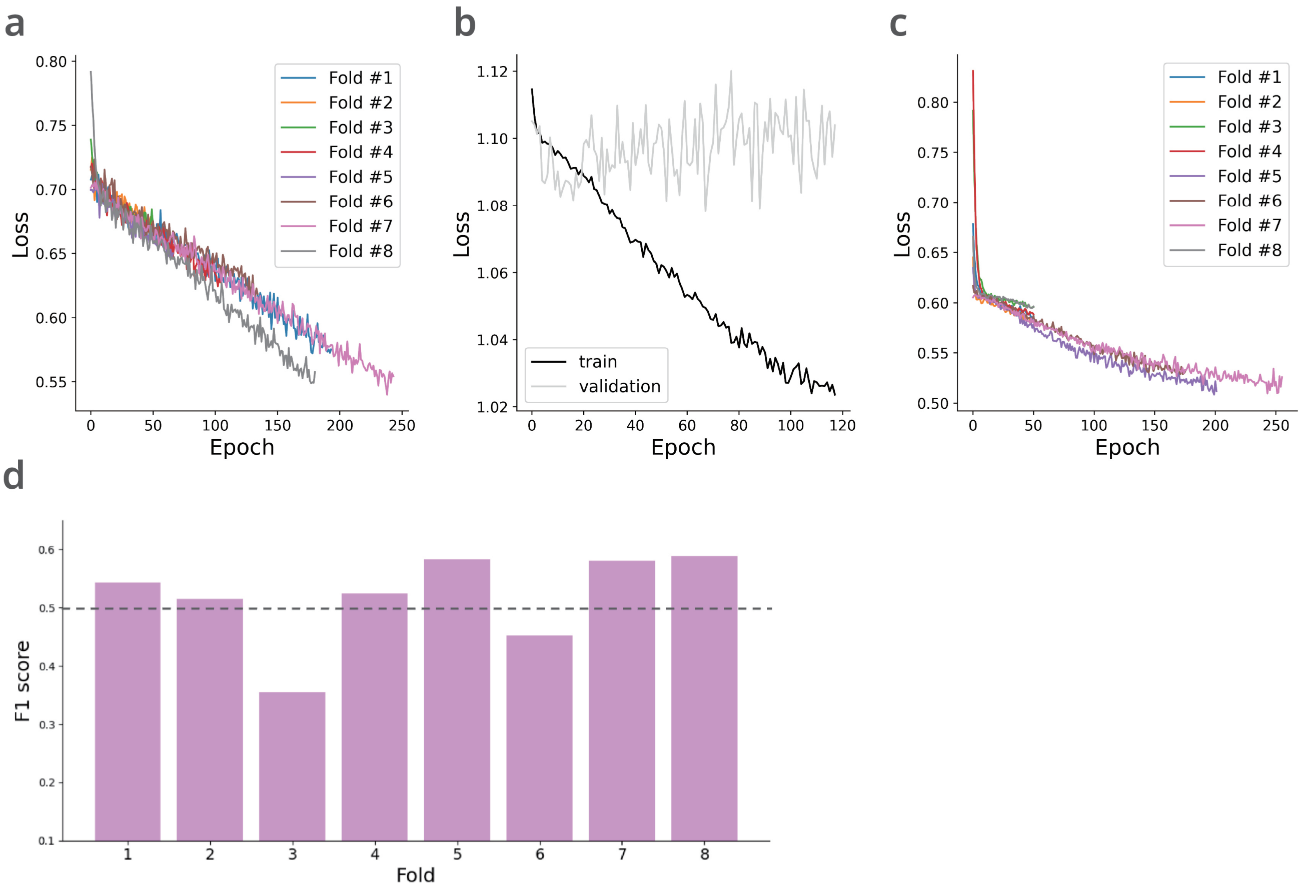


**Figure S3.** Training and performance details for the EEGNet classifier of a typical subject (Sub-06). (a) Training loss curve of the EEG-based model using the training-from-scratch approach. (b) Training loss curve of the EEG-based group model during the pre-training phase of the transfer learning approach. (c) Training loss curve of the EEG-based model during the fine-tuning phase of the transfer-learning approach (LvsO). (d) Macro F1 scores on the test set for each fold in cross-validation (Transfer Learning, LvsO). The dashed line indicates chance-level performance

| Subject | Accuracy | Precision | Recall | F1 score |
|---|---|---|---|---|
| sub-06 | 0.67 | 0.66 | 0.66 | 0.66 |
| sub-09 | 0.48 | 0.47 | 0.47 | 0.47 |
| sub-15 | 0.63 | 0.59 | 0.60 | 0.59 |
| sub-16 | 0.52 | 0.49 | 0.49 | 0.49 |
| sub-17 | 0.62 | 0.60 | 0.60 | 0.60 |
| sub-20 | 0.52 | 0.52 | 0.52 | 0.52 |
| sub-21 | 0.63 | 0.64 | 0.64 | 0.63 |
| sub-23 | 0.53 | 0.53 | 0.53 | 0.52 |
| sub-24 | 0.75 | 0.59 | 0.58 | 0.58 |

**Table S3.** Precision, recall, accuracy, and F1 score for the final ensemble model, aggregated across all subjects.

| Subject | Accuracy | Precision | Recall | F1 score |
|---|---|---|---|---|
| sub-06 | 0.55 | 0.54 | 0.54 | 0.54 |
| sub-09 | 0.55 | 0.53 | 0.53 | 0.53 |
| sub-15 | 0.66 | 0.61 | 0.61 | 0.61 |
| sub-16 | 0.53 | 0.51 | 0.52 | 0.51 |
| sub-17 | 0.54 | 0.53 | 0.53 | 0.53 |
| sub-20 | 0.52 | 0.52 | 0.52 | 0.52 |
| sub-21 | 0.66 | 0.69 | 0.68 | 0.66 |
| sub-23 | 0.52 | 0.53 | 0.53 | 0.52 |
| sub-24 | 0.68 | 0.46 | 0.47 | 0.47 |

**Table S4.** Precision, recall, accuracy, and F1 score for the final EEG-based model, aggregated across all subjects.

| Subject | Accuracy | Precision | Recall | F1 score |
|---|---|---|---|---|
| sub-06 | 0.66 | 0.64 | 0.65 | 0.64 |
| sub-09 | 0.49 | 0.52 | 0.52 | 0.49 |
| sub-15 | 0.61 | 0.60 | 0.61 | 0.59 |
| sub-16 | 0.53 | 0.51 | 0.51 | 0.51 |
| sub-17 | 0.54 | 0.53 | 0.53 | 0.53 |
| sub-20 | 0.51 | 0.51 | 0.51 | 0.51 |
| sub-21 | 0.50 | 0.50 | 0.50 | 0.50 |
| sub-23 | 0.52 | 0.52 | 0.52 | 0.52 |
| sub-24 | 0.61 | 0.56 | 0.60 | 0.54 |

**Table S5.** Precision, recall, accuracy, and F1 score for the final pose-based model, aggregated across all subjects.

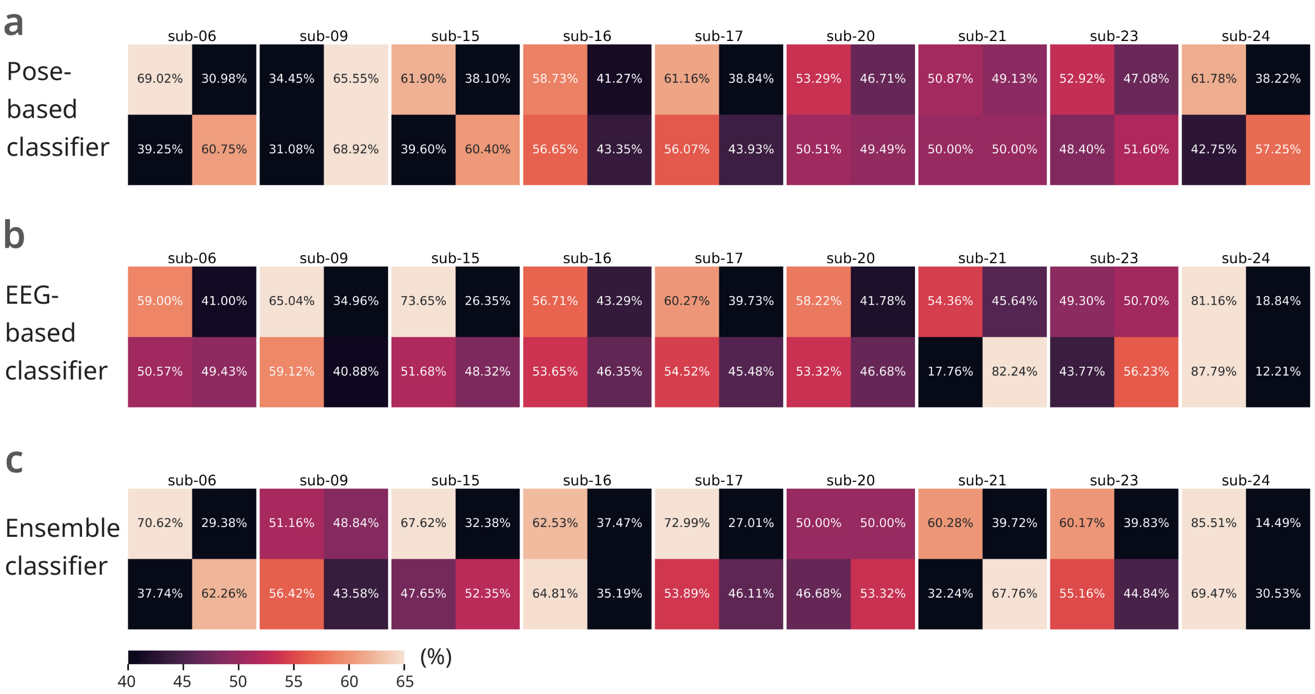


**Figure S4.** Per-subject confusion matrices (in %) for the (a) pose-based classifier, (b) EEG-based classifier, and (c) final ensemble classifier. Values represent mean classification percentages across subjects, with rows normalized by the true class (Left vs Right).

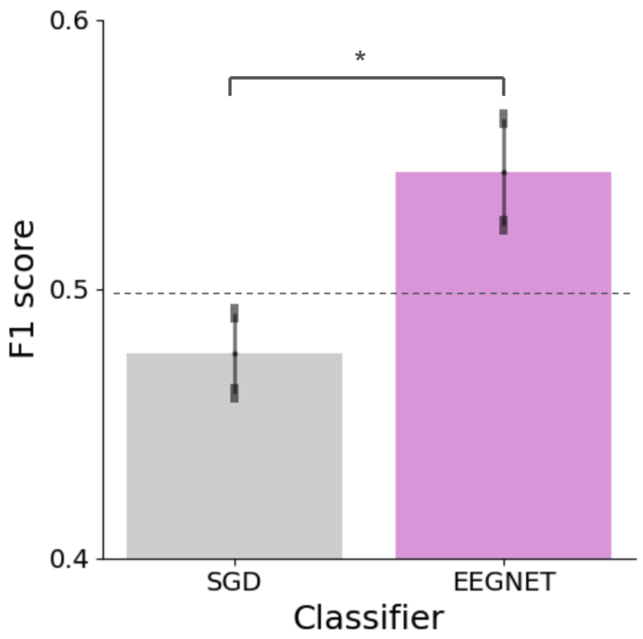


**Figure S5.** Statistical comparison of performance between ElasticNet and EEGNet models used as an EEG classifier at $-500$ ms before impact (paired t-test, $t(8) = 2.09$, $*p < 0.05$). The dashed line indicates chance-level performance. Error bars represent the SEM.

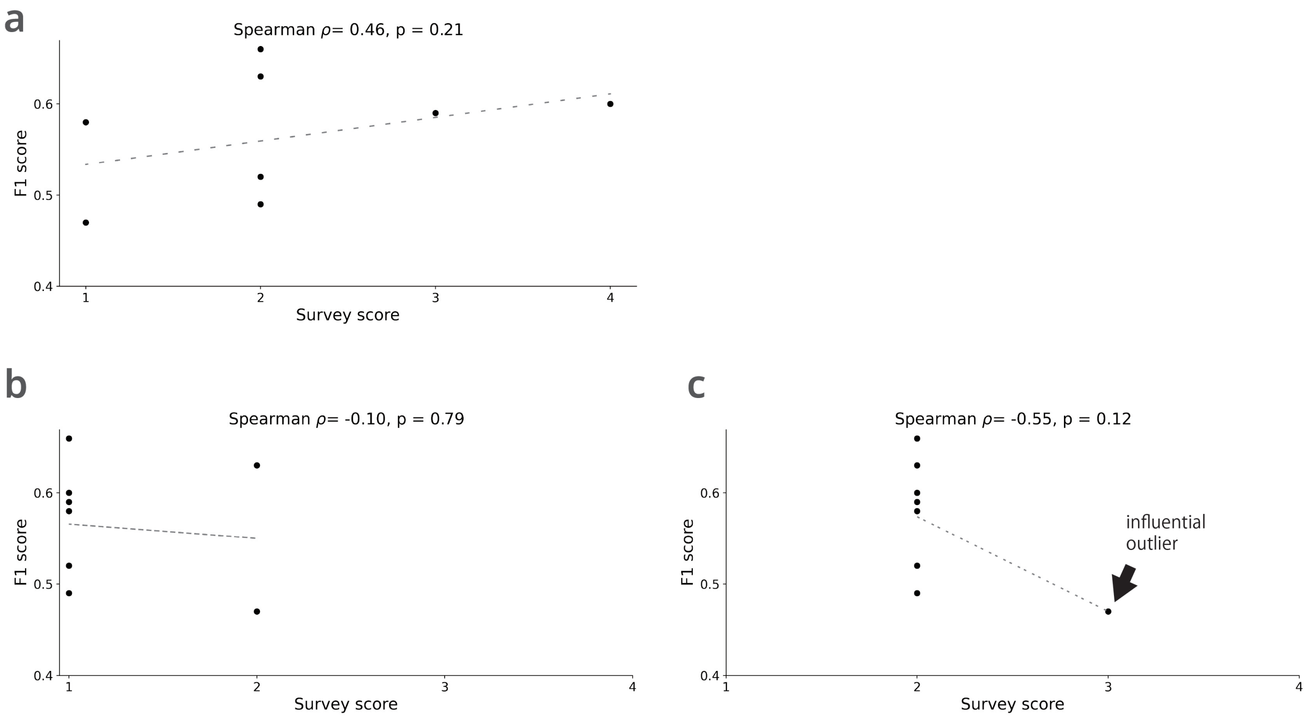


**Figure S6.** Association between ensemble model performance (macro F1 score) and (a) playing frequency (Q1), (b) tournament history (Q2), and (c) self-level assessment (Q3). Dashed lines indicate fitted monotonic trend lines. The black arrow indicates an influential outlier. Survey responses were encoded as ordinal values (1–4), with higher values indicating greater experience or proficiency.
